# Characterizing the targets of transcription regulators by aggregating ChIP-seq and perturbation expression data sets

**DOI:** 10.1101/2022.08.30.505909

**Authors:** Alexander Morin, Eric Ching-Pan Chu, Aman Sharma, Alex Adrian-Hamazaki, Paul Pavlidis

## Abstract

Mapping the gene targets of chromatin-associated transcription regulators (TRs) is a major goal of genomics research. ChIP-seq of TRs and experiments that perturb a TR and measure the differential abundance of gene transcripts are a primary means by which direct relationships are tested on a genomic scale. It has been reported that there is poor overlap in the evidence across gene regulation strategies, emphasizing the need for integrating results from multiple experiments. While research consortia interested in gene regulation have produced a valuable trove of high-quality data, there is an even greater volume of TR-specific data throughout the literature. In this study, we demonstrate a workflow for the identification, uniform processing, and aggregation of ChIP-seq and TR perturbation experiments for the ultimate purpose of ranking human and mouse TR-target interactions. Focusing on an initial set of eight regulators (ASCL1, HES1, MECP2, MEF2C, NEUROD1, PAX6, RUNX1, and TCF4), we identified 497 experiments suitable for analysis. We used this corpus to examine data concordance, to identify systematic patterns of the two data types, and to identify putative orthologous interactions between human and mouse. We build upon commonly used strategies to forward a procedure for aggregating and combining these two genomic methodologies, assessing these rankings against independent literature-curated evidence. Beyond a framework extensible to other TRs, our work also provides empirically ranked TR-target listings, as well as transparent experiment-level gene summaries for community use.

## Introduction

Understanding the regulatory interactions underlying gene expression programs is of considerable interest to contemporary experimental and computational biology. A fundamental objective is to map the relationships between transcription regulators (TRs) and the sets of gene targets they functionally influence. TRs, which include DNA sequence-specific transcription factors and chromatin proteins like MECP2 that bind methylated DNA, are a large class of proteins generalized by their ability to promote or repress gene activity (Lambert et al., 2018; Serebreni and Stark 2021). Learning the regulatory range of TRs is essential to understanding development, the functional identities of cell types, and the origins of diseases (Arendt et al., 2016, Lambert et al., 2018). However, experimentally establishing TR-target interactions is laborious and expensive, especially for precious tissues like the human brain. Additionally, efforts to predictively model these interactions as networks remains a challenging task complicated by a lack of known interactions (Marbach et al., 2012; Rothenberg, 2019; Nord and West, 2020). Identifying high-confidence sets of experimentally supported regulatory relationships is beneficial to inform biology as well as predictive method optimization.

We recently curated the literature for low-throughput biochemical assays demonstrating TR-target regulation (Chu et al., 2021), focusing on neurologically relevant TRs in mouse and human to expand upon existing resources like TRRUST (Han et al., 2018). These biochemical assays can provide strong evidence for direct regulation, but their coverage is understandably limited relative to the potential number of TR-target interactions. Accordingly, there are currently multiple genome-scale assays that provide regulatory information, but many rely on inference to determine TR-target relationships (Hawe et al., 2019). The most prominent means for high-throughput experimental assessment of direct interactions are to sequence purified DNA bound by an immunologically-selected TR (ChIP-seq), or to measure changes in gene transcript levels upon perturbation of the TR (differential expression – DE).

ChIP-seq and TR perturbation each have biological and technical considerations that complicate the task of assigning direct targets (Cusanovich et al., 2014; Kang et al., 2020). Consequently, applying both methods to a TR and intersecting the resulting gene lists is a common approach to enrich for regulatory interactions. This can be as simple as binarizing the significantly DE genes affiliated with a proximal ChIP-seq binding event, or through moderately more advanced strategies like the BETA algorithm, which combines the two gene lists into a single ordered ranking (Wang et al., 2013). However, efforts to evaluate the intersection of individual experimental pairs between these two genomic strategies in yeast (Hu et al., 2007; Gitter et al., 2009; Kang et al., 2020) and human (Cusanovich et al., 2014) have revealed that their evidence rarely converges.

Despite these complexities, ChIP-seq and perturbation experiments remain important strategies for biological discovery that continue to proliferate (Luo et al., 2020). Importantly, these data sets (or their intersection) are often used as the “gold standard” for evaluating computational strategies that predict TR-target interactions (Marbach et al., 2016; Miraldi et al., 2019; Pearl et al., 2019; Qin et al., 2020). In particular, two recent resources for human TRs have demonstrated the importance of aggregating distinct lines of evidence for the purpose of gene target assignment (Garcia-Alonso et al., 2019; Keenan et al., 2019). However, both focused solely on human TRs and held out the perturbation data to be used as a benchmark for their gene target rankings. Given the relative abundance of mouse gene regulation data, there is a clear need for the parallel organization and analysis of human and mouse TR-target interactions.

Here we set out a framework to identify and rank experimentally derived TR-target relationships by aggregating ChIP-seq and perturbation data sets from both mouse and human. We describe the degree of similarity across experiments and note characteristics of each data type that can complicate target assignment. We provide a framework for the aggregation and integration of these data sets, evaluating these empirically derived rankings against independent experimental evidence. This study also demonstrates how the collected information can potentiate further work, such as aligning bound regions to orthogonal genomic annotations, or identifying TR-target interactions with cross-species evidence.

## Results

For the current study we focused on an initial set of eight TRs to establish methodology and to calibrate expectations: ASCL1, HES1, MECP2, MEF2C, NEUROD1, PAX6, RUNX1, and TCF4 (not to be confused with TCF7L2, which is sometimes referred to as TCF4 in the literature). The selection of TRs was largely guided by our prior work on the curation of low-throughout experiments of neurologically relevant TRs (Chu et al., 2021). We emphasize that the data used in the current study were not required to be conducted in a brain-relevant system – our goal was to identify all ChIP-seq and high-throughput TR perturbation expression datasets for these regulators. Experiments excluded from analysis are noted in Supplemental Table S3. Our workflow is outlined in Fig. 1. In the subsequent sections, we give an overview of each genomic strategy before describing the aggregation and intersection approaches and their evaluation, leading to a consolidated ranking of candidate regulatory targets.

**Figure 1.**
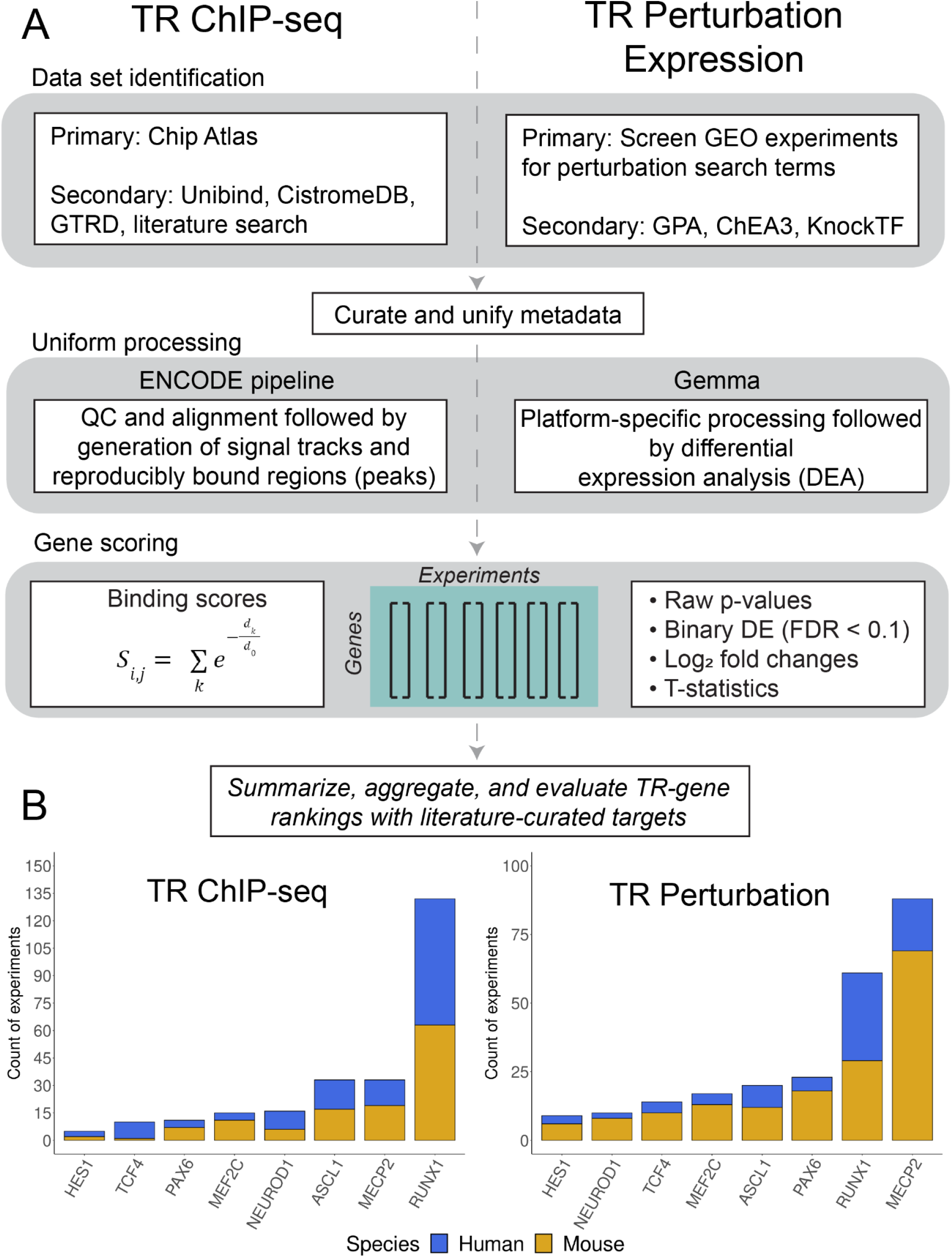
Study overview. (A) Workflow for TR ChIP-seq (left) and TR perturbation (right) data. (B) Counts of TR experiments considered for analysis.

### Identification, summarization, and gene-scoring ChIP-seq data sets

ChIP-seq data was predominantly identified across existing resources (particularly Chip Atlas – Zou et al., 2022) and supplemented with literature curation (Methods, Fig. 1A). All samples were curated into experimental units (based upon sample replication and presence of input controls) and uniformly processed using the ENCODE pipeline (Landt et al., 2012). A total of 255 experiments from 363 samples and 244 input controls were kept for analysis. While there was approximately equal representation of experiments across species, this equality does not extend to individual regulators (Fig. 1B; Supplemental Table S1).

Consistent with a previous effort to identify literature-sourced ChIP-seq data (Marinov et al., 2014), we found appreciable heterogeneity in the structure of experimental designs. Factors like the presence of an input control or sequencing depth can introduce technical variation to the count of inferred bound regions (peaks) in an experiment (Supplemental Fig. S1; Landt et al., 2012). This is an unavoidable reality when aggregating literature-sourced data, motivating our use of the stringent approach for peak calling promoted by ENCODE. Peaks were assigned to genes using a continuous scoring metric (from here referred to as the binding score; see Methods), and the ChIP-seq data were thus represented and analyzed as gene-by-experiment matrices of binding scores.

### ChIP-seq experiments targeting the same TR show moderately elevated similarity

As we aimed to aggregate TR data generated across distinct contexts, we first wanted to explore the similarity of binding profiles between the same TRs (intra-TR) and different TRs (inter-TR) across experiments. We examined both the Pearson correlation (*r*) of binding scores, which provides a measurement of similarity across all protein coding genes, as well multiple measures of overlap for the top 500 scoring genes (Sikora-Wohlfeld et al., 2013; Keenan et al., 2019).

The results were consistent regardless of the approach, showing that, collectively, intra-TR experiments were moderately more similar than inter-TR pairs (Figs. 2A-C; Supplemental Fig. S2). Specifically, on average intra-TR data sets shared 57/500 top genes, while inter-TR pairs shared 22/500. As expected, the most similar experiments were from the same GEO submission (done by the same lab), followed by experiments conducted in comparable biological contexts by distinct research groups. For example, the highest global correlation (*r* = 0.87, 308/500 top scoring genes) was between two different ASCL1 constructs in the SH-SY5Y neuroblastoma cell line (GSE153823: Ali et al., 2020), while the most correlated experiments from distinct groups both assayed Ascl1 in the developing mouse neural tube (*r* = 0.66; 251/500 top scoring genes; GSE43159: Sun et al., 2013; GSE55840: Borromeo et al., 2014).

**Figure 2.**
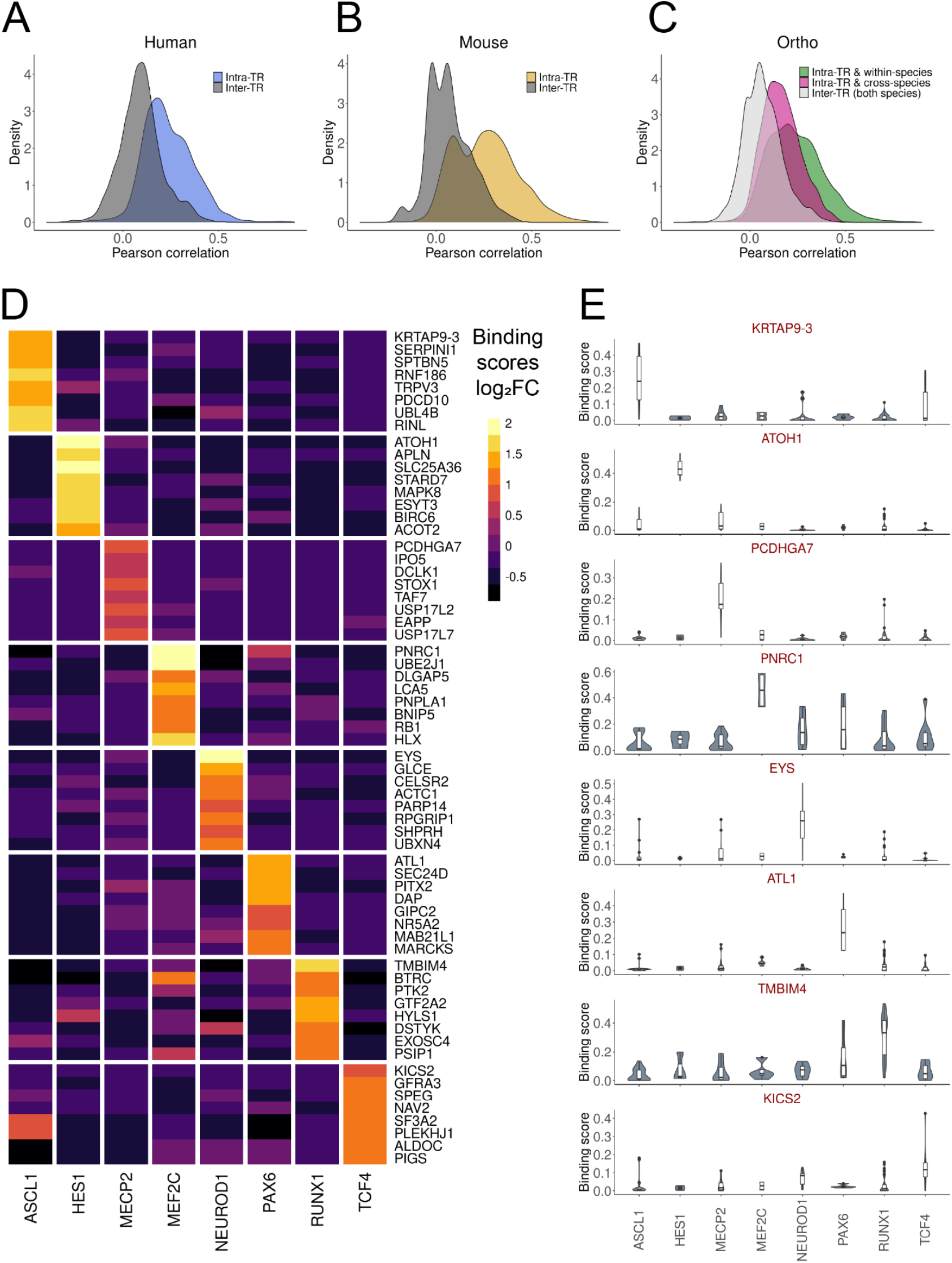
ChIP-seq experiment similarity and specifically bound genes. (A-C) Distribution of binding score correlations between ChIP-seq experiments targeting the same TR (intra-TR) versus different TRs (inter-TRs). (D) Log_2_ fold changes of binding scores for the top eight ranked genes (by p-value) for each TR in human experiments using a mixed effect linear model. As scoring was distance-based, linearly proximal genes had similar ranks, and so for plotting only the most significant genes (*PCDHGA7* for MECP2 and *KRTAP9-3* for ASCL1) are shown. (E) Distributions of binding scores for the most significant gene for each TR from the same model as in (D).

Elevated intra-TR correlation was not universal to all comparisons (Supplemental Fig. S2), which in some instances may be attributable to cell-type patterns. For example, the three human HES1 experiments had intra-correlations ranging from *r* = 0.16-0.19. A HES1 experiment from this trio conducted in the K562 cell line had inter-correlations ranging from 0.29-0.36 with five other K562 experiments targeting NEUROD1 or RUNX1. However, all intra-HES1 pairs had more genes in their top 500 overlap (range 69-92) than any inter-HES1 comparison (range 2-63). We also found instances where there was less intra-TR similarity than might be expected. Two ENCODE K562 RUNX1 experiments had an *r* = 0.32 (69/500 top scoring), despite targeting the same TR in the same cell type. These experiments used different RUNX1 antibodies and sequence library strategies, serving as a reminder of the considerable technical variation of the ChIP-seq methodology.

Using the same approach, we compared the similarity of mouse and human ChIP-seq experiments, based on orthology of TRs and of targets. For targets, we focused on a set of 16,976 high-confidence one-to-one mouse-human orthologs (Methods). The distribution of binding score correlations for the same TR but different species (intra-TR & cross-species) was shifted higher relative to both inter-TR & cross-species and inter-TF & within-species comparisons (collapsed as one group for simplicity in Fig. 2C; Supplemental Fig. S2C). The top of these intra-TR & cross-species rankings were dominated by RUNX1 experiments: of the 2,351 intra-TR & cross-species comparisons that had greater than average common top scoring genes (>37/500), 2,089 (90%) were associated with RUNX1. This is attributable to the relatively high abundance of data for this TR, and the frequency in which RUNX1 experiments were conducted in blood contexts in both species, consistent with conserved regulatory profiles leading to cellular identities (Arendt et al., 2016). While slightly attenuated, the shift in correlation distributions was held when excluding RUNX1 experiments (Supplemental Fig. S3).

Collectively, these observations support that consistent TR binding profiles may be identified across studies, albeit with an expected loss of highly context-dependent signals. Despite this potential for false negatives, this was nevertheless promising for our goal of aggregating data to uncover consistent evidence in support of specific TR-target relationships.

### A mixed effect linear modeling framework identifies genes with TR-enriched binding

Finding evidence for intra-TR binding similarities, we looked to identify and rank the bound genes for each TR. While we ultimately used the intra-TR mean binding score in our final aggregated rankings (Discussion), we found that certain regions had a propensity to be bound generically, consistent with prior observations (Supplemental Fig. S5; Discussion). For example, the constitutively expressed *GPAA1* was found to have a peak within 25 kbp of its transcription start site (TSS) in 104/129 (81%) of the human experiments, distributed across all TRs. We therefore developed a strategy to identify candidate TR targets that could address the concern of binding specificity (Methods). Briefly, we used a mixed effect linear modeling framework with TR identity as the main effect, accounting for high-level experimental factors and the heightened correlation among experiments generated by the same group. While not without caveats (Discussion), this approach was designed to seek evidence for selective TR binding patterns despite the heterogeneity of contexts typically found in each comparator group.

Fig. 2D demonstrates the results of this approach, plotting the differential binding scores of the eight most significant genes for each human TR, while 2E shows the boxplots of the binding scores of each TR’s most significant gene (mouse in Supplemental Fig. S4). This yielded an average of 643 candidate target genes per TR (log_2_ fold change (FC) > 0 & false discovery rate (FDR) < 0.05), ranging from three (mouse Hes1) to 1,398 (human RUNX1). This model revealed a number of previously characterized TR-target interactions. Well-described ASCL1 targets and Notch pathway effectors *DLL1, DLL3, DLL4,* and *HES6* (Castro et al., 2006; Nelson et al., 2009; Castro et al., 2011) had elevated binding in both human and mouse ASCL1 comparisons, while the HES1 target *ATOH1* (Kazanjian and Shroyer, 2011) was the most significant gene in the human HES1 comparison. Taken together, this analysis supported that aggregated ChIP-seq data can prioritize independently described regulatory interactions. We more formally evaluate this approach in a later section.

### Identifying frequently bound loci relative to regulatory element annotations

The framework described thus far is gene-centric, relying on a scoring metric that sums the contribution of binding events around a TSS. Consequently, a gene may have a high binding score for a given TR, even if the individual binding events are dispersed around the TSS across experiments. We therefore identified the regions most commonly bound by a TR, providing discrete coordinates for future investigation (Supplemental Data S5).

To demonstrate an application of these bound loci, we examined their overlap with candidate cis-regulatory elements (cCREs) (Moore et al., 2020). These regions, which encompass predicted promoter and enhancer-like regions, were previously defined through the integration of multiple genomic features across diverse cellular contexts. Genomic annotations like the cCREs aim to characterize the biological function of the underlying DNA sequences. To provide context before focusing on frequently bound loci, we first examined the proportion of each ChIP-seq experiment’s peaks that overlapped each class of cCRE (Fig. 3).

**Figure 3.**
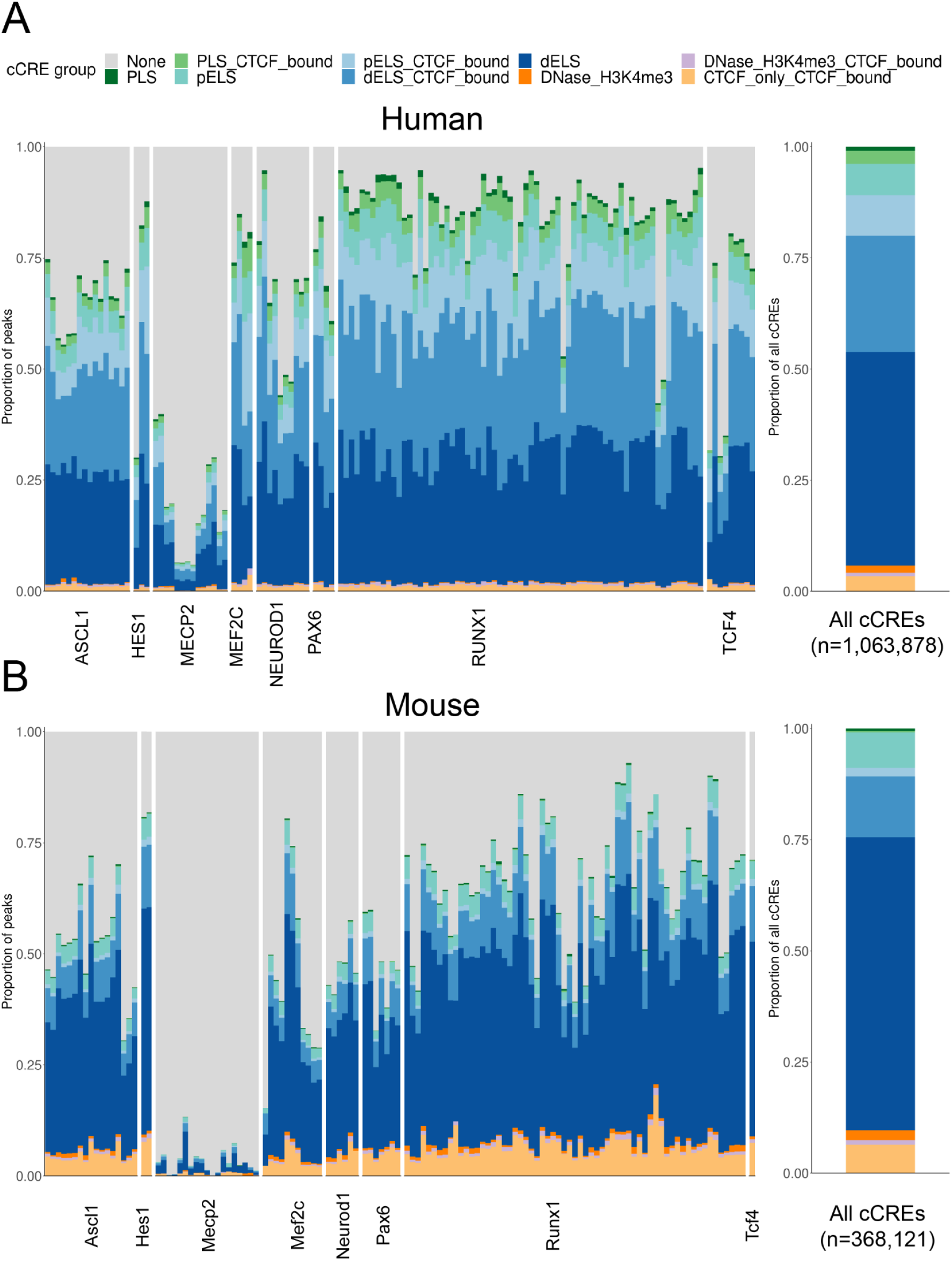
Overlap of ChIP-seq peaks with annotated regulatory elements. (A) Left panel: The proportion of peaks for each human ChIP-seq experiment, grouped by TR, that overlapped with a candidate cis-regulatory element (cCRE; PLS = promoter-like sequence; ELS = enhancer-like sequence; p = TSS proximal; d = TSS distal). Right panel: The proportional breakdown of all cCRE groups. (B) Same as in (A), except for mouse ChIP-seq experiments.

With the exception of MECP2, we found that 74% of the peaks in a typical human ChIP-seq experiment overlap with a cCRE (Fig. 3A), and in mouse 59% (Fig. 3B). We note that mouse cCRE groups were defined with less input data than human and are expected to have fewer discovered elements and thus less overlap. And while there was variation across TF experiments in how peaks were distributed across the cCRE groups, they generally followed the global cCRE group proportions, with most binding to regions characterized as enhancer-like sequences. MECP2 was the exception: the overlap of peaks dropped to an average of 16% for human and 4% for mouse, even though MECP2 experiments had a comparable number of peaks to the other experiments (Supplemental Fig. S1C). This lack of overlap may reflect MECP2’s differential mode of binding relative to the TFs (Shah and Bird 2017).

As discussed earlier, the concordance of binding across experiments for a TR was limited, but by focusing on areas of agreement we hope to uncover meaningful biology. In the locus-specific analysis, there are many cases of highly reproducible sites. For example, for human ASCL1 we found 29 regions bound across all 16 experiments, and 405 regions bound in at least 14, many of which were infrequently bound by the other TRs (Supplemental Fig. S6). A notable example is a ~330 bp sequence in an intronic region of *SHB*, annotated as an enhancer-like cCRE, which had a peak called in all ASCL1 experiments but only twice in non-ASCL1 experiments (2/113).

We take such patterns of reproducibility and specificity as an indication of biological relevance. However, not all binding, even if reproducible, is expected to result in significant regulatory activity (Wasserman and Sandelin 2004; Teytelman et al., 2013; Cusanovich et al., 2014), hence the importance of considering orthogonal data (Garcia-Alonso et al., 2019). This brings us to the introduction of the TR perturbation experiments, which provides evidence of TR regulation at the RNA level. For example, the candidate ASCL1 target *SHB* we identified in the above binding analysis is differentially expressed (DE) in 6/8 human ASCL1 perturbation experiments, raising our confidence in its relevance. In the next sections, we describe the systematic analysis of TR perturbation experiments for integration with the ChIP data.

### Acquisition and summarization of TR perturbation data sets

TR perturbation data was predominantly identified using a screen of GEO experiments and supplemented with existing resources (Methods, Fig. 1A). All were processed in the Gemma database (Lim et al., 2021), and a total of 242 experiments were considered for downstream analysis (Supplemental Table S2). Sequencing platforms were slightly more represented than microarrays (Supplemental Fig. S7). Unlike the ChIP-seq collection, where there were similar amounts of mouse and human data, the TR perturbation corpus was heavily skewed towards mouse, with just over twice as many experiments in mouse than human (Fig. 1C), although the breakdown by TR was broadly similar. Gene knockouts were the most common perturbation strategy, making up nearly half of all experiments (Supplemental Fig. S7). Overexpression and knockdowns followed with near equal representation. The remainder we classified as “mutants”, following the original authors’ descriptions. These are somewhat distinct from knockouts, typically (but not always) involving loss of function-inducing point mutations rather than larger deletions.

### TR perturbation experiments show modest differential expression effect sizes

We first explored the properties of the collected perturbation data prior to any aggregation. Consistent with Cusanovich et al. (2014), we found that the perturbation effect sizes tended to be modest. Given the distribution of fold changes (FC) across all experiments (Fig. 4A), we did not apply FC thresholding and classified genes as differentially expressed (DEGs) at a relaxed FDR < 0.1. This framework resulted in a median count of 261 DEGs per experiment (adding a constraint of a minimum absolute FC of 1 would result in a median of 34).

**Figure 4.**
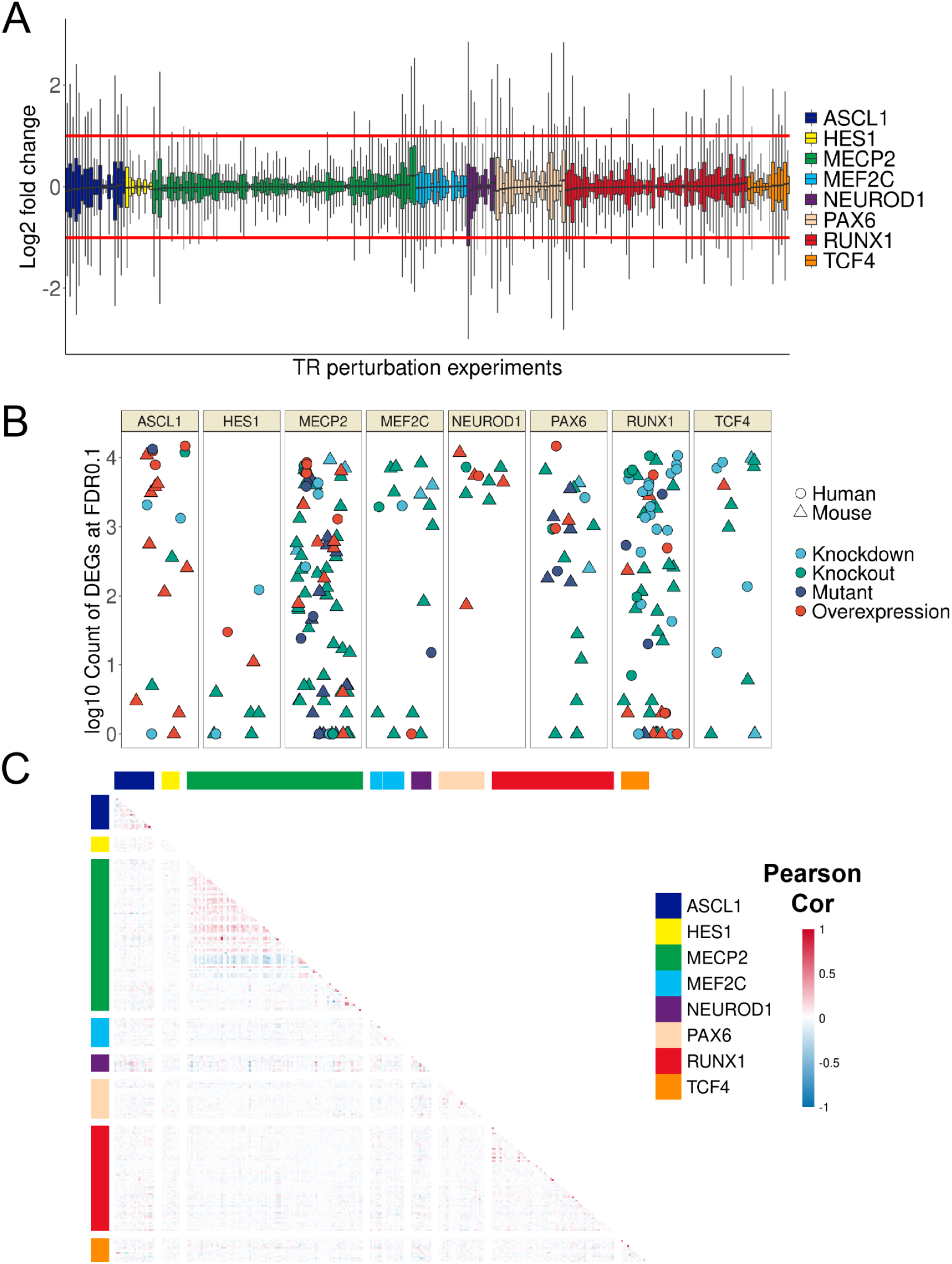
Overview of TR perturbation experiments. (A) Distribution of gene log_2_ fold changes (FC) across the 242 mouse and human perturbation experiments. FC was clipped at [−3, 3] for plotting. (B) Count (log_10_ scale) of differentially expressed genes grouped by TR; colour denotes perturbation strategy and shape denotes species. (C) Heatmap of correlation values of gene FC between experiments, grouped by TR. Note that only orthologous genes were calculated here to allow plotting of both species; the relatively minimal intra-TR correlation holds when considering mouse and human separately.

Also in line with Cusanovich et al., we found that the number of genes affected by a TR perturbation was poorly predicted by the FC magnitude of the perturbed TR (Supplemental Figs. S9B,C). For example, there was an appreciable number of experiments that had no DEGs despite substantial changes in the TR’s expression level. Otherwise, there was a spread in DEG counts for each TR, with NEUROD1 having the most on average and HES1 the least (Fig. 4B). While biological characteristics may explain these extremes (e.g., pioneering activity of NEUROD1 versus a more repressive role for HES1), the variety of designs, contexts, and sample sizes complicate these comparisons. Still, we noted a difference in the count of DEGs associated with perturbation type, with knockdowns having the highest median (1,950) and knockouts the lowest (112) (Supplemental Figs. S10A,B).

### Despite select examples, intra-TR perturbation experiments show weak similarity

As with the ChIP-seq collection, we explored the similarity of intra-TR versus inter-TR perturbation experiments. Because there was unequal gene coverage across experiments due to platform differences (Supplemental Fig. S7), we calculated Pearson correlations of log_2_ FCs (as well as their absolute values) for the genes commonly measured between pairs of experiments. For the top 500 overlap comparison, we separately considered up- and down-regulated genes, as well as sorting by p-values from the DE analysis.

Consistent with expectations, the most similar experiments were intra-TR comparisons from the same research group, led by a pair of ASCL1 overexpression experiments (r = 0.95, 410/500 by up-regulated; GSE153823: Ali et al., 2020). As for ChIP-seq, experiments from different groups but comparable contexts showed elevated similarity, such as two experiments overexpressing Ascl1 in astrocyte-to-neuron conversions (*r* = 0.60, 222/500 overlap of up-regulated genes; GSE174238: Kempf et al., 2021; GSE132674: Rao et al., 2021). We also found examples of anti-correlative patterns that aligned with the opposing roles of the perturbations, typically involving a *MECP2* overexpression versus a *MECP2* loss-of-function (e.g., *r* = −0.66, 49/500 by p-value; GSE126640: Cholewa-Waclaw et al., 2019).

While these examples are somewhat encouraging, intra-TR similarities as a group were only marginally different from inter-TR comparisons (Fig. 4C; Supplemental Fig. S8). Focusing on the top 500 overlaps by p-values, an average human intra-TR pair shared 32/500 genes to the 28/500 of inter-TR pairs; mouse intra-TR pairs had 29/500 compared to 21/500 in inter-TR pairs. These trends extended to the orthologous gene comparison between species. The strongest correlation was between *NEUROD1* overexpression studies in mouse and human that both looked to generate neuronal populations (*r* = 0.32, 135/500 by up-regulated; GSE104435: Matsuda et al., 2019; GSE149599: Pomeshchik et al., 2020), while the strongest negative correlation was between a human *MECP2* overexpression in a neuronal cell line and a mouse *Mecp2* knockout in cortical neurons (*r* = −0.24, 39/500 by p-value; GSE126640: Cholewa-Waclaw et al., 2019; GSE124521: Keidar et al., 2019). However, a typical intra-TR & inter-species comparison shared only three more genes in a top 500 comparison relative to the average inter-TR pair.

### Aggregating perturbation data sets reveals repeatedly DE genes

We next used an aggregation approach to identify consistent patterns across perturbation experiments for each of the TRs. This is in keeping with our overall philosophy of taking advantage of commonalities while being cognizant of the issues noted above. To rank genes for each TR, we used a simple tally of the count of times a gene was DE across intra-TR experiments (Count DE), breaking ties with the average absolute FC.

Encouragingly, we identified many genes that were frequently DE for a given TR’s set of perturbation experiments (Fig. 5C). The exception was HES1, which had less data overall and had few DEGs in both species (Fig. 4B). Mouse Mecp2 had the most perturbation data and correspondingly had the genes with the highest Count DE, led by *Irak1* with 24/69 in mouse Mecp2 experiments and 5/19 in human. This NF-κB pathway gene has previously been associated with Mecp2 (Urdinguio et al., 2008; Kishi et al., 2016; Supplemental Note). However, we were unable to find literature support for other highly ranked Mecp2 candidates, such as the brain-enriched estrogen-related receptor *Esrrg* (20/69 mouse, 7/19 human) which also had strong Mecp2 binding scores in our ChIP-seq analysis, suggesting that further investigation into this interaction is warranted.

**Figure 5.**
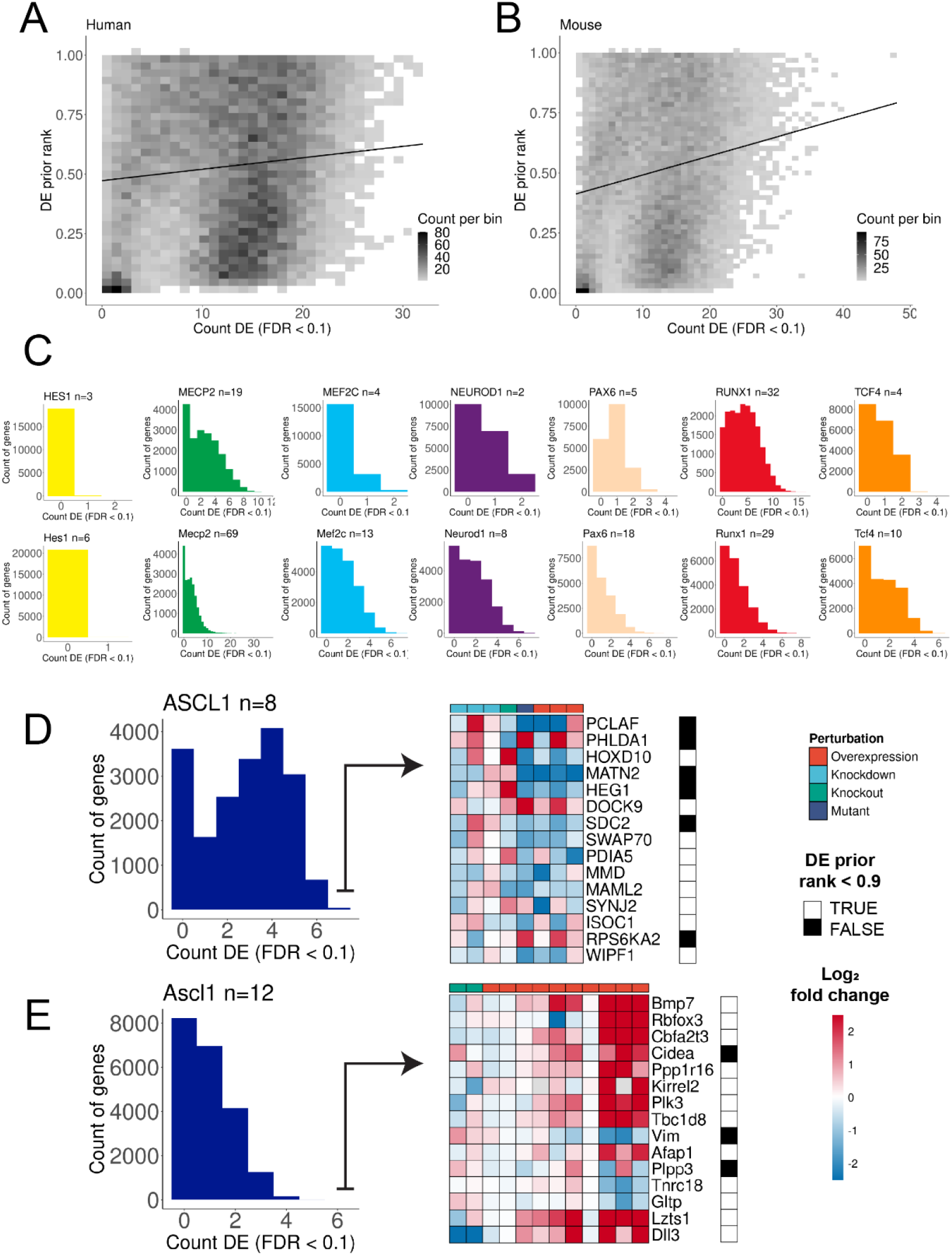
Demonstration of genes with recurrent differential expression (DE). (A) The x-axis is the amount of times that a gene was DE across human experiments in the current study (n=77) and the y-axis is the DE prior rank, where 1.0 represents the gene that was most likely found as DE across a large and diverse corpus of expression experiments. (B) Same as in (A) but for mouse experiments (n=165). (C) Histograms of the count of times genes were DE across each group of TR experiments (upper row human, lower row mouse). (D) Demonstration of the top 15 genes by DE evidence for human ASCL1. FCs are clipped at [−2.5, 2.5] for plotting. The DE prior was binarized so that values above 0.9 (black squares) represent genes that are commonly DE regardless of design. (E) Same as in (D) but for mouse Ascl1.

We also observed many orthologous genes with recurring DE in both species, such as the neurogenic growth factor *BMP7* for ASCL1 (6/8 human, 6/12 mouse). Human NEUROD1 had the fewest perturbation experiments (n = 2), yet we identified five genes that were DE in both human studies as well as in seven of eight for mouse Neurod1 (9/10 of all NEUROD1 experiments): *PTPRK, UGCG, SFT2D2, SLC35F1,* and *SOGA1*. *SLC35F1* was recently identified as an under-studied neurodevelopmental gene implicated in epileptic encephalopathies (Di Fede et al., 2021); our work thus connects NEUROD1 to the regulation of this presumed synaptic plasticity gene. On the other hand, there were also many examples of orthologous genes that were repeatedly measured as DE in one species but never in the other - such as MECP2 candidates *SLC6A7* (15/69 mouse, 0/19 human) or the pro-neural *NRG2* (0/69 mouse, 8/19 human) - but ruling these as species-specific targets requires further consideration beyond the scope of this study.

### Additional considerations of perturbation data aggregation

We highlight two final considerations of aggregating the perturbation data. First, akin to the “generic” signals observed in the ChIP-seq collection, we observed genes that were frequently DE across TR studies (Supplemental Figs. S10B,C and S11). We compared these counts with a previous metric that ranks genes by their predictability of being DE (DE prior; Crow et al., 2019), finding a weak but significant trend in both human (*r* = 0.11, p-value < 2.2e-16) and mouse (*r* = 0.18, p-value < 2.2e-16) (Figs. 5A,B). This trend could reflect biological underpinnings of the observations of Crow et al., but the weakness of the signal precluded making strong conclusions. Second, we observed genes could have discordant directions of change (up- vs down-expressed) across the same type of perturbation for a given TR. We quantified this by adapting the metric of *Purity* (Methods), scoring the consistency of a gene’s FC direction across loss- and gain- of function experiments (Supplemental Fig. S12). As we do not assume that frequently DE genes are not true targets, and that a given TR can be activating or repressive in different contexts (Lambert et al., 2018), we elected not to incorporate *Purity* or the DE prior into our final rankings. Still, we believe these metrics provide additional context, such as for identifying candidate interactions that are predominantly activating or repressive.

### Combining aggregated ChIP-seq and perturbation evidence to prioritize gene targets

Having summarized each line of genomics evidence, we turned to our original goal of ranking gene targets by the combined evidence of the two strategies. We first simply examined the degree of overlap in their gene lists. Given the generally weak perturbation intra-TR experiment similarity, it was perhaps unsurprising to see the same trend when comparing across ChIP-seq and perturbation experiment pairs (Supplemental Fig. S13). A typical intra-TR ChIP-seq and perturbation experiment pair shared 20/500 top-ranked genes versus 17/500 for inter-TR pairs.

However, much like the perturbation comparisons, we observed many genes with frequent intra-TR overlaps between the two methods, despite the overall weak group overlap. To reach a consolidated list for each TR, we extended a popular rank product approach previously used on individual ChIP-seq and perturbation experiment pairs (Breitling et al., 2004; Tang et al., 2011; Wang et al., 2013; Methods). Rank products have been used extensively in genomics and other fields, representing a simplistic yet robust meta-analytic summarization of noisy data (Koziol et al., 2010). Here, we used the aggregated perturbation and ChIP-seq gene lists as inputs, rather than individual experiments, to avoid biases for TRs with imbalanced data set counts between the two methods. Thus, for each TR, we obtained aggregated gene orderings for both genomic lines of evidence and a final combined ranking (integrated).

To evaluate the integrated rankings, we wished to use an orthogonal line of experimental evidence for comparison, similar to a recent benchmark of human TF-target interactions that used perturbation data as the gold standard (Garcia-Alonso et al., 2019). In contrast to Garcia-Alonso et al., we cannot use perturbation data as an evaluation set as we used it to generate the rankings. We reasoned that the information from TR perturbation data was more important to use for gene prioritization rather than to hold it out for assessment, particularly given that low overlap was already an expectation given our observations and prior work (Garcia-Alonso et al., 2019; Kang et al., 2020). We thus turned to resources that curated low-throughput interactions (Chu et al., 2021; Supplemental Note). These yielded 483 unique targets for the eight TRs, ranging from 11 for TCF4 to 156 for PAX6 (median 51; Supplemental Fig. S16). Importantly, this collection is not exhaustive, lacks annotation of negatives (non-target genes), and is diverse in contexts, so we do not consider it a true gold standard. However, we reasoned that it would still provide at least a sense of whether the integrated genomics data could help prioritize known (and thus, unknown) interactions, relative to the performance of individual data sets or a single data modality.

First, we tested the difference in the aggregated gene rankings by presence in the curated set (Supplemental Figs. S14, S15). All mouse rankings showed evidence for prioritizing curated targets (Wilcoxon test p-value < 0.05), save for the Tcf4 binding and Neurod1 perturbation aggregations. This potentially may be explained by the size of the evaluation set, as these two TRs had the fewest curated targets (Supplemental Fig. S16). Notably, some curated Neurod1 targets like *Insm1* (Breslin et al., 2003) were not highly ranked (perturbation rank 793rd) despite being DE in 4/8 mouse Neurod1 experiments. Thus it is possible that the genes ranked higher than *Insm1* by DE evidence include numerous real but unidentified targets. For example, *Nova2* is a neurodevelopmental splicing regulator (Mattioli et al., 2020) that was DE 6/8 times and had enriched Neurod1 binding, suggesting this may be a true interaction lacking low-throughput evidence. While the human results were more mixed – none of the three MEF2C differential tests were significant, unlike mouse Mef2c – the majority of human comparisons still provided evidence that the rankings prioritized curated targets. In sum, this analysis supported that our aggregation strategy was able to assign heightened importance to genes with independent experimental evidence.

### Aggregate rankings typically outperform null expectations and single experiments

The previous analysis considered the difference in median ranks by curation status. To more directly assess the ability of the aggregation strategies to preferentially rank known interactions, we performed a precision-recall analysis, calculating the area under the precision-recall curve (AUPRC) to summarize performance (Marbach et al., 2012; Fig. 6 and Supplemental Fig. S16). We first note the overall low performance of each of these rankings (e.g., ASCL1 in Fig. 6B), an unsurprising result given factors like the incomplete nature of the evaluation set (Discussion). The integrated rankings achieved a higher AUPRC than the single method aggregations for human ASCL1, mouse and human RUNX1, mouse Pax6 and human TCF4. However, the individual data type aggregations also sometimes outperformed the integrated ranking.

**Figure 6.**
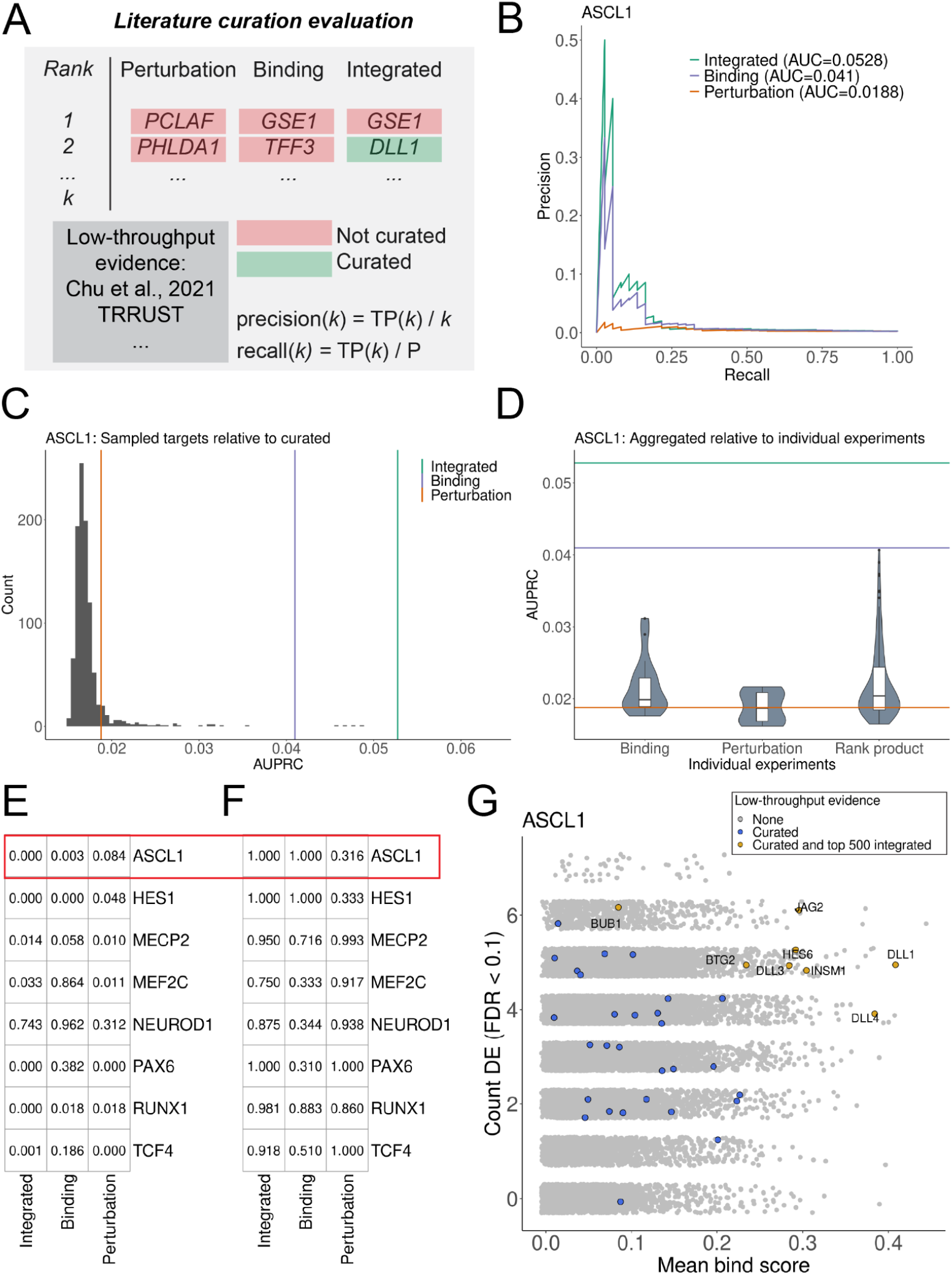
Overview of literature curation evaluation framework. (A) Precision and recall were calculated at every step (*k*) of the aggregated rankings for presence in the curated target resource. TP = true positives called at step *k*, P = all curated targets for the current TR. (B) PR curve and the associated areas under the curves (AUPRC) for human ASCL1. (C) The distribution of AUPRCs when sampling random targets from the curated resource and using the integrated ASCL1 ranking to calculate the AUPRC. Bars refer to the observed performance when using the curated ASCL1 targets. (D) Distribution of AUPRCs when using the individual ASCL1 experiments or their rank product pairings to order genes. (E) Proportion of samples in whose AUPRC exceeded the observed values. (F) The percentile of the aggregate AUPRCs relative to the distribution of all individual comparisons in (D). (G) Example of ASCL1 targets with genomics and low-throughput evidence.

To better contextualize these relative differences in performance, we conducted two further comparisons using the precision-recall framework. First, we created a null distribution of AUPRCs for each TR, iteratively sampling curated targets from the entire literature resource and calculating the AUPRC with the aggregated rankings and sampled targets. This analysis revealed that, despite the low overall performances, almost every TR had an aggregate ranking that exceeded the null. Figs. 6C,E demonstrates that no null target set outperformed the human ASCL1 integrated rankings, a trend also seen for rankings like mouse Pax6 and human RUNX1. Further, the human MEF2C perturbation and integrated rankings outperformed the null expectation, despite the lack of MEF2C differential ranking by curation status (Supplemental Figs. S14, S16). Thus, while aggregation for lesser-represented TRs may assign curated targets a similar rank to uncurated targets *on average*, the top of the aggregated ranking can still be enriched for curated targets relative to a null expectation.

Finally, we compared the aggregated AUPRCs to those obtained when calculated from individual ChIP-seq and perturbation experiments, or their individual rank-product pairings. This provided a direct comparison of the performance of the aggregated rankings to what is obtained from the individual experiments constituting the aggregation. Consistent with the previous analyses, most TRs had an aggregated ranking whose AUPRC exceeded the expected values from individual experiments, even if this was not always the integrated aggregation (Figs. 6D,F and Supplemental Fig. S16). The human ASCL1 integrated ranking, for example, outperformed every individual ASCL1 experiment, as well as the perturbation and binding aggregations, supporting the benefit of both data aggregation and cross-method integration.

Similarly, both the human and mouse RUNX1 integrated rankings were in the 98th percentile of RUNX1 AUPRCs, outperforming the perturbation and binding aggregations. While the integrated ranking also outperformed the vast majority of individual experiments, the top human RUNX1 performances were typically rank-product pairings conducted in leukemic systems. As the most represented RUNX1 cell types in the curated resource were leukemia cell lines, the benchmark may be biased towards cell-type-specific effects compared to the cell-type agnostic aggregations. This may also explain why mouse Neurod1 had the least performant aggregations: the highest AUPRCs belonged to individual genomic experiments (and their rank-product pairings) conducted in pancreatic tissues. In concordance, pancreatic experiments were among the most represented from the relatively scarce collection of literature curated NEUROD1 targets, while the genomic NEUROD1 experiments covered a broader range of tissues.

In conclusion, while the integrated rankings were not always the most performant, the aggregation strategies presented are typically still more capable of prioritizing known targets than null expectations or the individual experiments composing the aggregation. All summarized rankings are found in Supplemental Data S1 and the contributing data in Supplemental Data S2.

## Discussion

In this study, we build upon existing strategies (Tang et al., 2011; Wang et al., 2013) to create a framework for aggregating genomics data to rank gene regulatory relationships. Crucially, we focused on methodologies that directly assess TR activity, rather than those based on inference (e.g., coexpression or DNA motif footprinting). Prior studies have similarly ranked TR-target interactions using orthogonal lines of evidence (Garcia-Alonso et al., 2019), or aggregated TR-target libraries (Keenan et al., 2019). While these studies are more comprehensive in their TR coverage, they focus only on human interactions, and in particular hold out TR perturbation evidence for benchmarking rather than for gene prioritization. Here, we did a detailed analysis of a subset of TRs to better understand the properties of the contributing genomics experiments and to identify potential biases of aggregation. Towards this end, our work also represents a novel meta-analysis of TRs with demonstrated importance in mammals. We discuss a subset of the identified candidate targets in the Supplement Note.

Data re-use and aggregation across diverse studies presents many challenges. Extending prior observations, we find generally weak similarities when comparing experiments targeting the same TR (Hu et al. 2007; Cusanovich et al. 2014; Garcia-Alonso et al., 2019; Kang et al., 2020). Further, the genomics evidence (aggregated or not) was not highly performant when evaluated using literature-curated targets. This is unsurprising, given (1) the incomplete nature of the evaluation set, (2) the heterogeneity of contexts and experimental/technical factors in all considered data, and (3) the inherent difficulty in benchmarking gene regulation (Marbach et al., 2012; Garcia-Alonso et al., 2019). The first reason in particular guided our use of the broadly applicable rank product for data integration (Breitling et al., 2004; Wang et al., 2013). Here, this is a two parameter model that gives equal importance to each genomic line of evidence, while training TR-specific models would require a robust gold standard for each TR. Nevertheless, our data aggregation revealed numerous candidate TR-target interactions supported by extensive convergent evidence. As we (generally) see that the aggregated data prioritized curated targets, our explicit assumption is that the rankings will also be enriched for unexplored direct interactions.

It is important to discuss caveats of our framework, as some of these complexities extend to general difficulties in studying gene regulation. First and foremost, our rankings must be interpreted as a sorting of existing genomics evidence, rather than one of absolute biological importance. If a developmentally-critical interaction is not assayed in the appropriate biological or temporal context of the included experiments, it cannot be expected to be highly ranked – if it can even be captured by the considered strategies. Similarly, a highly context-specific interaction will not be as highly prioritized as one common to a more abundantly represented context, due to our approach of aggregating data. Therefore, researchers interested in benchmarking predictions may wish to use the aggregated rankings, while those interested in a specific context may focus on the targets identified by a subset of the assembled data deemed most relevant (Supplemental Data S2).

For the binding evidence, our gene scoring method aligns with other contemporary studies analyzing ChIP-seq data at scale (Methods). While providing more granularity than binarizing binding events, these formulations still rely on genomic distance as a measure of relevance to a gene (Chen et al., 2020). Current evidence suggests this to be a useful approximation (Yoshida et al., 2019), but future efforts may benefit from incorporating evidence from 3D chromosomal interactions, such as in the “activity by contact” model (Fulco et al., 2019). Additionally, the TSS-based logic of the binding score is likely better suited for TFs than other classes of regulators like MECP2. Although we were still able to capture curated MECP2 targets, researchers may choose to prioritize the perturbation rankings or focus on specific MECP2-bound coordinates (Supplemental Data S3-S5).

In line with prior reports, we also find that certain loci are frequently associated with a ChIP-seq signal across assays (Supplemental Fig. S5). The biological-versus-technical nature of these regions has been debated (Teytelman et al., 2013; Wreczycka et al., 2019; Partridge et al., 2020; Ramaker et al., 2020); regardless of its origin this phenomenon motivated our analysis of differential binding activities (Fig. 2). For example, Li et al., (2019) subtracted a gene-wise background signal when gene scoring a single ChIP-seq experiment. However, our study was based on a small and biased selection of TRs, thus the comparator groups may participate in common regulatory pathways. For this reason, we based the final binding rankings on the mean binding score, rather than on differential binding statistics. Nevertheless, we find evidence for specific binding that aligns with prior described interactions (e.g., HES1-ATOH1, ASCL1-DLL1). We believe that the binding score-based linear modeling framework has intriguing potential for forming more sophisticated TR group comparisons, such as for co-binding partners or TR families.

For the perturbation evidence we tallied independent significance tests. Finding that genes commonly had variable changes in FC direction, we ultimately used the absolute FC as a tie-break. Together, this means that the final rankings are agnostic to the change of direction, although our inclusion of FC *Purity* in the summarized results allows researchers to identify interactions predominantly measured as activating or repressive. Ideally a single model would jointly consider all TR perturbation experiments. However, this is greatly complicated by the diversity of technologies, gene coverage (non-uniformity of missing observations), sample sizes, and designs, thus we elected for simplicity in this study. We used the DE prior (Figs. 5A,B) to provide additional context of a gene’s behavior with respect to DE testing, as it can help identify “generically” DE genes.

Genes with frequent DE evidence showed a range of binding evidence across TRs. Although our primary interest was finding targets corroborated by both strategies, genes well-supported by TR perturbation alone may warrant further investigation. These possibly reflect instances where the assembled ChIP-seq contexts did not capture the binding event, or that it occurred at a genomic distance missed by our scoring scheme. Alternatively, a common interpretation for individual experiment pairs is that DE genes lacking in binding are indirectly regulated through an intermediate regulator. If an absence of binding can be confirmed it is strongly suggestive that, even if indirect, the perturbed TR and frequently DE gene participate in a tightly controlled regulatory pathway.

While TRs are expected to have cell-type specific targets, our results nominate genes that may be regulated by a TR across contexts (Gertz et al., 2013; Lambert et al., 2018). While an ultimate goal of gene regulation research is to establish the specificity of TR-target interactions, it would still be desirable to characterize the extent by which each TR can be represented by a set of “core” targets. ASCL1, for example, appears to commonly regulate Notch pathway effectors like *DLL1, DLL3,* and *HES6* (Castro et al., 2006; Nelson et al., 2009; Castro et al., 2011). Further examination of these frequent targets is warranted: if the same regions are bound or if there is more distributed enhancer usage; the degree to which these interactions are coexpressed across systems/conditions; conservation of the associated sequence, and the consistency of chromatin features or co-binding partners. Our examination of frequently bound loci is a step in this direction (Supplemental Fig. S6), but a more comprehensive exploration building on these observations requires further study. Similarly, a more comprehensive examination of TR-target interactions conserved across species is warranted, which can be potentiated by the examples provided by our work.

Consistent with our low-throughput curation resource (Chu et al., 2021), we found that the literature coverage of the studied regulators was greatly uneven, with MECP2 and RUNX1 receiving the most attention in both low-throughput and high-throughput investigations. This isn’t surprising, but highlights the need for investigation of less-studied yet important regulators. Similarly, we identified multiple examples of genes that had convergent lines of evidence but are sparsely represented in the literature (Supplemental Note). Given that the assembled data is “biased” towards regulators deemed of interest by the broader research community, our work suggests that these under-studied candidate targets could be prioritized by gene functionality studies.

In sum, we believe this study will be a useful resource for researchers interested in gene regulation. We present a large collection of transparently summarized information that catalogues the current state of the literature, while also potentiating novel biological discovery.

We also have documented many practical issues and limitations of the considered data and present an analytical framework that is readily extensible to the ever-growing collection of TR experimentation.

## Methods

Except where noted, analyses were performed in the R statistical computing environment (R Core Team 2022).

### Genomic feature tables

Gene annotations were based on NCBI RefSeq Select (mm10 and hg38), which assigns one TSS to each gene (https://www.ncbi.nlm.nih.gov/refseq/refseq_select/). ENCODE candidate cis-regulatory elements (cCREs) were obtained from https://screen.encodeproject.org/ (V3; (Moore et al., 2020). The DE prior rank information was an updated version for human and newly generated for mouse, using the same strategy as in Crow et al., 2019 but expanded to a greater number of expression platforms and data sets (Supplemental Tables S4, S5). High-confidence one-to-one orthologous genes were accessed via the DIOPT resource (V8; Hu et al., 2011), keeping only genes with a score of at least 5 that were also reciprocally the best score between mouse and human, and excluding genes with more than one match.

### Identification of ChIP-seq data sets

Identification of ChIP-seq data was predominantly facilitated using the Chip Atlas database (Zou et al., 2022) owing to its breadth of mouse and human data and the organization of the associated metadata. Additional experiments were identified in the literature and other ChIP-seq resources: GTRD (Kolmykov et al., 2021), Unibind (Puig et al., 2021), and Cistromedb (Zheng et al., 2018). Experiments were curated and matched to their input controls, applying and extending Chip Atlas’s metadata framework such that each row corresponds to a unique *SRX* ID (https://www.ncbi.nlm.nih.gov/sra).

### Uniform processing of ChIP-seq data

As there was heterogeneity across each resource’s processing pipelines and how sets of samples were organized within an experiment unit (e.g., the pooling of replicates or inputs), we uniformly re-processed all data. Using *SRX* IDs, sample library information was obtained using NCBI’s *esearch* utility (version 13.8) and fastq files were downloaded using *fasterq-dump* (version 2.10.8) before read trimming and quality control were performed using *Trim Galore* (version 0.6.6), keeping reads that were at least 30 bp. Processed fastq files were then submitted for processing in the comparatively stringent ENCODE ChIP-seq pipeline (version 1.3.6) (Landt et al., 2012; https://github.com/ENCODE-DCC/chip-seq-pipeline2) with the following considerations: First, samples were grouped into units based upon the replicate status of the experimental design, with all identified input controls pooled for each experimental unit. We note that many experiments consisted of a single replicate (meaning that the reproducibility process operated on pseudoreplicates rather than true replicates), and that input controls could be shared across distinct experimental units. Second, we fixed the peak caller to *MACS2*, as the ENCODE default method *SPP* cannot be used for runs with no input controls. Finally, all samples were submitted to the TF pipeline procedure, save for MECP2 experiments, which were submitted to both the TF and histone procedures. Given MECP2’s binding profile, prior studies have used broad peak-calling parameterization (Gabel et al., 2015; Ito-Ishida et al., 2018; Xiang et al., 2020). Correspondingly, more MECP2 experiments succeeded processing with the histone parameterization, and thus we used this strategy for all MECP2 samples. We followed the advice of Marinov et al., 2014 to avoid applying flat QC cut-offs for heterogeneous ChIP-seq collections, with two exceptions: Experiments marked as “fail” in the reproducibility analysis (IDR for TF, Overlap for MECP2) in the generated QC report were excluded, and only experiments with at least 100 peaks were retained to avoid overly sparse binding vectors during analysis. We further note that these excluded experiments typically also had outlier ENCODE QC metrics.

### ChIP-seq gene binding scores and normalization

The ENCODE pipeline produces a set of output files, among which is a single “optimal reproducible” peak set table of genomic coordinates (via the IDR procedure for TF and Overlap procedure for histone/MECP2) for each experimental unit - we focused on these tables for most analyses. We considered multiple approaches to score gene binding for each experiment. The most common strategy is binary assignment, where genes are scored as 1 if a peak summit (the most read-enriched base pair) is found within a distance threshold to the TSS, and 0 otherwise. Following the advice of Sikora-Wohlfeld et al., 2013, we focused on quantitative binding scores, in particular a slight modification to the exponential decay function introduced in Ouyang et al., 2009:

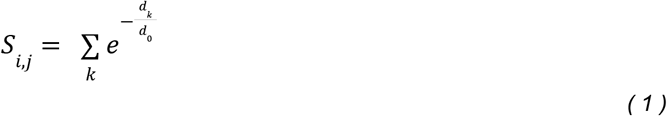

Where S is the binding score, *i* indexes a protein coding gene’s TSS, *j* indexes a TR experiment, *k* represents all peak summits within 1Mbp of the current TSS, *d_k_* represents the absolute distance in bps between the 5’ TSS and the peak summit, and d_0_ is the decay constant, set to 5,000 as in the original publication. The original formulation scaled each element by the *MACS2* intensity score, which we omitted as these scores no longer retained their original interpretation after the ENCODE reproducibility process. The omission of this scaling factor is consistent with other work that adopt an exponential decay formulation to score genes using ChIP-seq peaks (Wang et al., 2013; Garcia-Alonso et al., 2019; Chen et al., 2020). Thus, all ChIP-seq experimental units can be represented as a gene by experiment matrix of binding scores. We added 1 and applied a log_10_ transformation followed by quantile normalization (*preprocessCore* R package version 1.48) to these bind score matrices, finding that this strategy helped alleviate batch/technical considerations, and used these matrices for the reported PCA analysis, mean bind scores, and ranking individual genes.

### Binding specificity analysis

To find genes with enriched binding scores for the same TR, we adopted the limma-voom framework (version 3.42.2; Law et al., 2014), a common strategy for applying linear models to genomics data with a positive mean-variance relationship, as observed here. The raw binding score matrices were submitted to the voom transformation with quantile normalization specified, and using limma’s *duplicateCorrelation* function with laboratory identity as a blocking variable to account for expected elevated correlations among experiments submitted by the same research group. The following model was then fit:

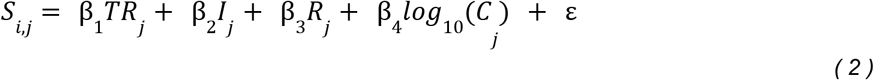

Where *S, i,* and *j* are the same as in equation 1, the main effect *TR* represents the ChIP’d protein, binary variables *I* and *R* capture if experiment *j* has at least one input control and at least one replicate, and *C* is the count of peaks for experiment *j*, with the residual (ε) having the covariance matrix as estimated by duplicateCorrelation, and regression weights provided by voom. Finally, for each TR, a “one-versus-rest” contrast was extracted from this model, which estimates for each gene the difference in mean binding scores (*S*) for the current TR’s set of experiments relative to all other experiments, after using information from all experiments to account for the specified experimental structure/technical variables.

### ChIP-seq peak region overlap

All overlap procedures were performed using the *GenomicRanges* R package (version 1.38; Lawrence et al., 2013). For the cCRE region analysis, only the peak summit was used to detect overlap across any part of a cCRE region (which ranged from 150-350 bps). For the frequently-bound region analysis, as the original peaks were variable in length, we re-sized each such that 150 bps was added in each direction from the summit.

### Identification of TF perturbation high-throughput expression data sets

Perturbation strategies can be coarsely grouped by whether they reduce the available pool of TR gene transcripts (knockdowns), if one or both of the TR alleles are functionally eliminated (knockouts), if a transgenic construct results in elevated levels of the TR (overexpression), or if sequence variations critically disrupt the function of the TR (mutants). We first queried existing resources that have aggregated TF perturbation experiments: Gene Perturbation Atlas (Xiao et al., 2015), ChEA3 (Keenan et al., 2019), and KnockTF (Feng et al., 2020). Most experiments were identified by extending strategies used by our group for the Gemma database (Lim et al., 2021). Briefly, this involves human curation of experiment suitability after programmatically searching the GEO database for co-occurrence of TR gene symbols and a list of perturbation terms (e.g., “siRNA”, “overexpression”). All identified experiments were checked for accurate curation by at least two individuals. Selected experiments were required to have at least two control and treatment samples, have samples that perturbed only the single TR of interest, and were of an appropriate technical strategy (single-cell sequencing, sorting by expression, and run-on sequencing were excluded).

Surprisingly, we found that multiple experiments showed minimal or even “unexpected” expression changes in the perturbed TF (e.g., overexpression of TR yielding an apparent decrease in RNA levels for the TR) (Supplemental Fig. S9A). We inspected all such examples, finding that the perturbed TR was often among the top ranked genes by absolute fold change (median 81^st^ percentile). While we cannot exclude the possibility of sample mislabeling on GEO, it is possible that temporal or biological factors such as proposed genetic compensatory mechanisms (El-Brolosy and Stainier 2017; El-Brolosy et al., 2019) may explain the measured TR transcript levels post-perturbation. Given that the associated studies typically validated the perturbation independent of the microarray or RNA-sequencing experiment, we chose to keep all such experiments.

### Obtaining summarized TR perturbation results from Gemma

The selected expression studies were submitted to the Gemma framework for uniform processing (Lim et al., 2021). This entails human curation of the experimental design using controlled terminology, paired with automated handling of batch information, platform-specific support, and differential expression analysis (DEA). Gemma fits generalized linear models using the curated experimental factors, producing a table of summarized results for each factor (t-statistics, log_2_ fold changes, and p-values). We note that some studies had experimental factors beyond the TF perturbation (e.g., “+/− LPS treatment”). When distinct cell types/tissues were one of these factors (e.g., a knock-out in hypothalamus as well as cerebellum), a separate perturbation DEA was performed for each cell type/tissue. Otherwise, a single model was fit for all experimental factors and the perturbation contrast effect sizes (controlling for the other factors) were extracted. Microarray probes that did not map to a single gene were excluded, and when multiple probes or sequencing elements mapped to a single gene, only the element with the maximum absolute t-statistic was kept. The false discovery rate was controlled using the Benjamini-Hochberg procedure (*p.adjust* R version 3.6.0) on the p-values after this filtering. Genes were binarized as differentially expressed (DE) at FDR < 0.1. In this manner, all experiments can be represented as gene by experiment matrices of the various effect sizes. Because microarrays vary in which genes are assayed, their inclusion resulted in variable gene coverage in the final corpus (Supplemental Fig. S7).

### Purity of TR-perturbation change of direction

To quantify whether a gene was predominantly up/down-regulated across the different types of perturbation experiments, we adopted the *Purity* metric used in clustering analysis to determine cluster quality. We treated gain-of-function (GoF – overexpressions) and loss-of-function (LoF – knockdowns and knockouts; mutants were excluded) experiments as two clusters in which genes are classified as either up-regulated (FC > 0) or down-regulated (FC < 0):

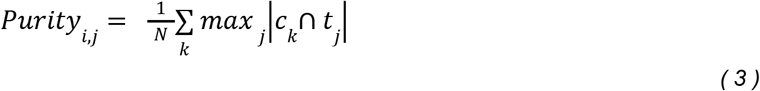

Where *i* indexes genes, *j* indexes TRs, N is the number of non-mutant experiments in which gene *i* is measured for TR *j*, *k* is the number of clusters (two – GoF and LoF), *c_k_* is a cluster, and *t_j_* is the dominant classification within that cluster. If gene *i* has FC < 0 in 3/4 GoF experiments (GoF is dominantly down-regulated) and FC < 0 in 4/4 LoF experiments (LoF is dominantly down-regulated) for TR *j*, then *Purity_i,j_* = (3 + 4) / 8 = 0.875. We further define *Signed purity*, which considers the dominant class/direction across both clusters: If the dominant direction is the same for GoF and LoF, we multiply *Purity* by −1 (thus the above example becomes −0.875), otherwise *Purity* is preserved. In this manner we can distinguish genes that show consistent changes in direction across experiments but unexpected convergence across the perturbation types. In this study, we arbitrarily define consistent genes as having *Purity* > 0.8.

### Ranking gene targets

For perturbation data we ranked genes by the count of times that they were DE across experiments (Count DE), using the absolute log_2_ fold change as a tie-break (*D*), and for ChIP-seq we used the mean binding score (*S)*. We considered multiple strategies to reach a single ranking from both data types. A popular approach for individual experiment pairs is the BETA algorithm, which takes the rank product (RP) of a DE gene list and binding scores from an exponential decay function based on distance from the TSS (as in this study) (Wang et al., 2013). However, we wished to rank targets using all the intra-TR data sets, not just pairs. Two options are to tally the count of times a gene was in a “top overlap,” or to average a gene’s RP across every intra-TR comparison. However, each requires making numerous non-independent comparisons (each experiment is paired multiple times). Alternatively, rank aggregation strategies have a long history of use in genomics (e.g., Keenan et al., 2019 averages rankings across different TR-target libraries), with strategies extended to cases involving unevenly sized rankings (Kolde et al., 2012; Li et al., 2019). Yet, the final ranking will be influenced if there are imbalanced experiment counts between the two genomic methods, and adjusting for this would require calibration against a known standard. Consequently, we used a simplistic hybrid approach, taking the rank product of the two aggregated lists:

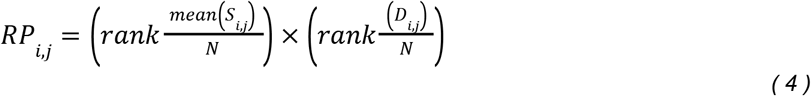

Where *N* is the number of genes. The RP returns a unitless value, which we in turn convert to a rank such that 1 represents the most prioritized gene *i* for TR *j*.

## Data access

The code used for analysis in this study can be found at https://github.com/PavlidisLab/TR_aggregation. The gene rankings and underlying data matrices used for analysis can be found as RDS objects in the Borealis data repository at https://doi.org/10.5683/SP3/MAFGFL.

## Acknowledgments

We thank Dr. Nathaniel Lim for the expansion of the DE prior ranking (manuscript in preparation), first generated in Crow et al. (2019). We also thank Dr. Marine Louarn, who has managed the update of the resource of curated regulatory interactions (Chu et al., 2021). This work was supported by National Institutes of Health grant MH111099 (https://www.nih.gov/) and Natural Sciences and Engineering Research Council of Canada grant RGPIN-2016-05991 (https://www.nserc-crsng.gc.ca/), both held by PP. The funders had no role in study design, data collection and analysis, decision to publish, or preparation of the manuscript. AM had funding support from CIHR-CGS, NSERC CREATE, and IMH Marshall Scholars programs.

