## Supplemental material for "Characterizing the targets of transcription regulators by aggregating ChIP-seq and perturbation expression data sets"

***Supplemental material table of contents***

*Supplemental Figures* ……………………………………………………………………………… *2*

*Low-throughput curated target resources* ……………………………………………………………*18*

*Overview of TR targets* …………………………………………………………………………….. *19*

*Supplemental citations* …………………………………………………………………………….. 27

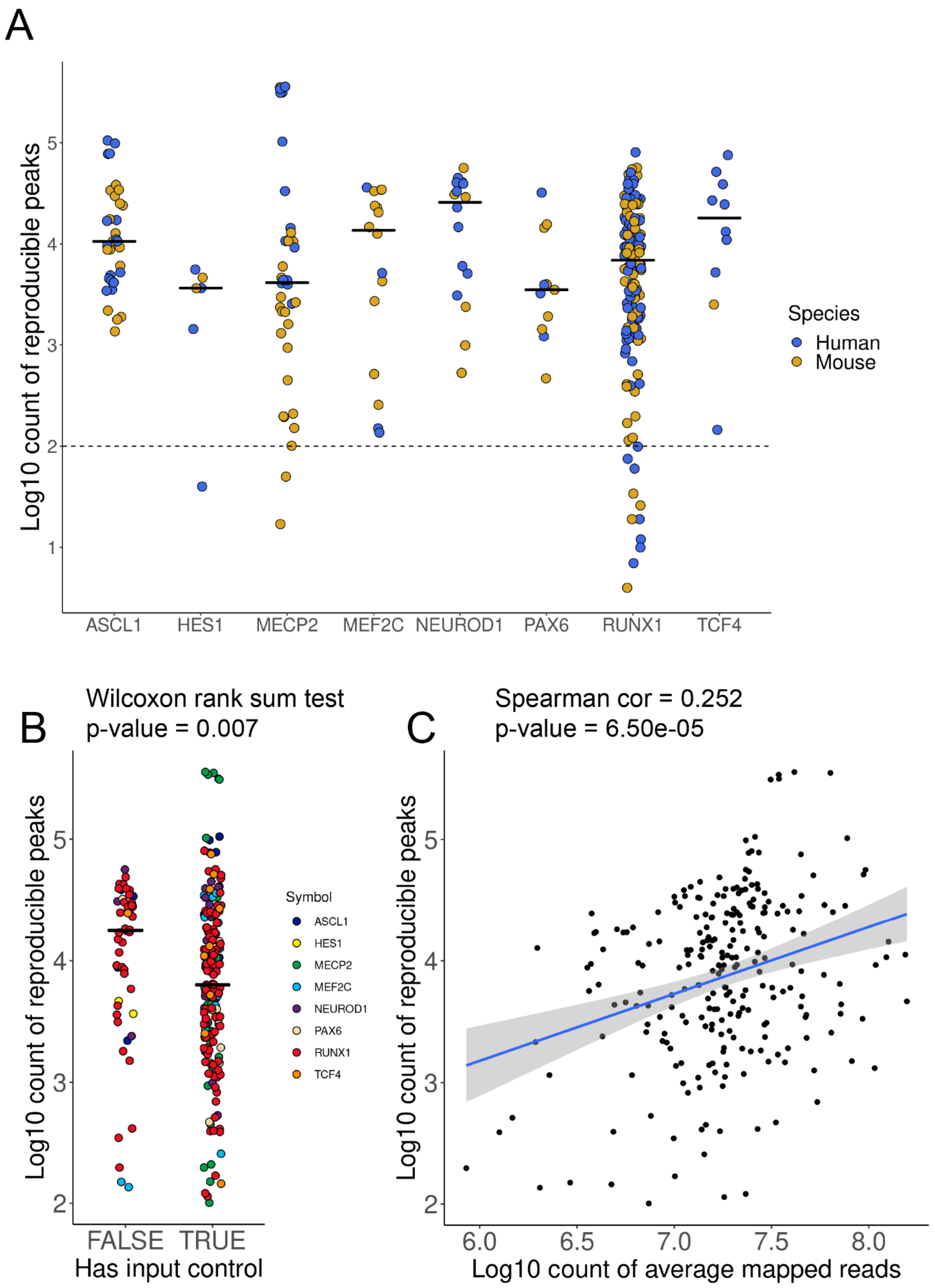


**Figure S1**. Overview of ChIP-seq experiment count of reproducible peaks. (A) Count of peaks by TR and coloured by species; dashed line represents minimum peaks required for analysis. (B) Count of peaks by presence of at least one input control. (C) Count of peaks by averaging de-duplicated and mapped reads across samples within an experimental unit.

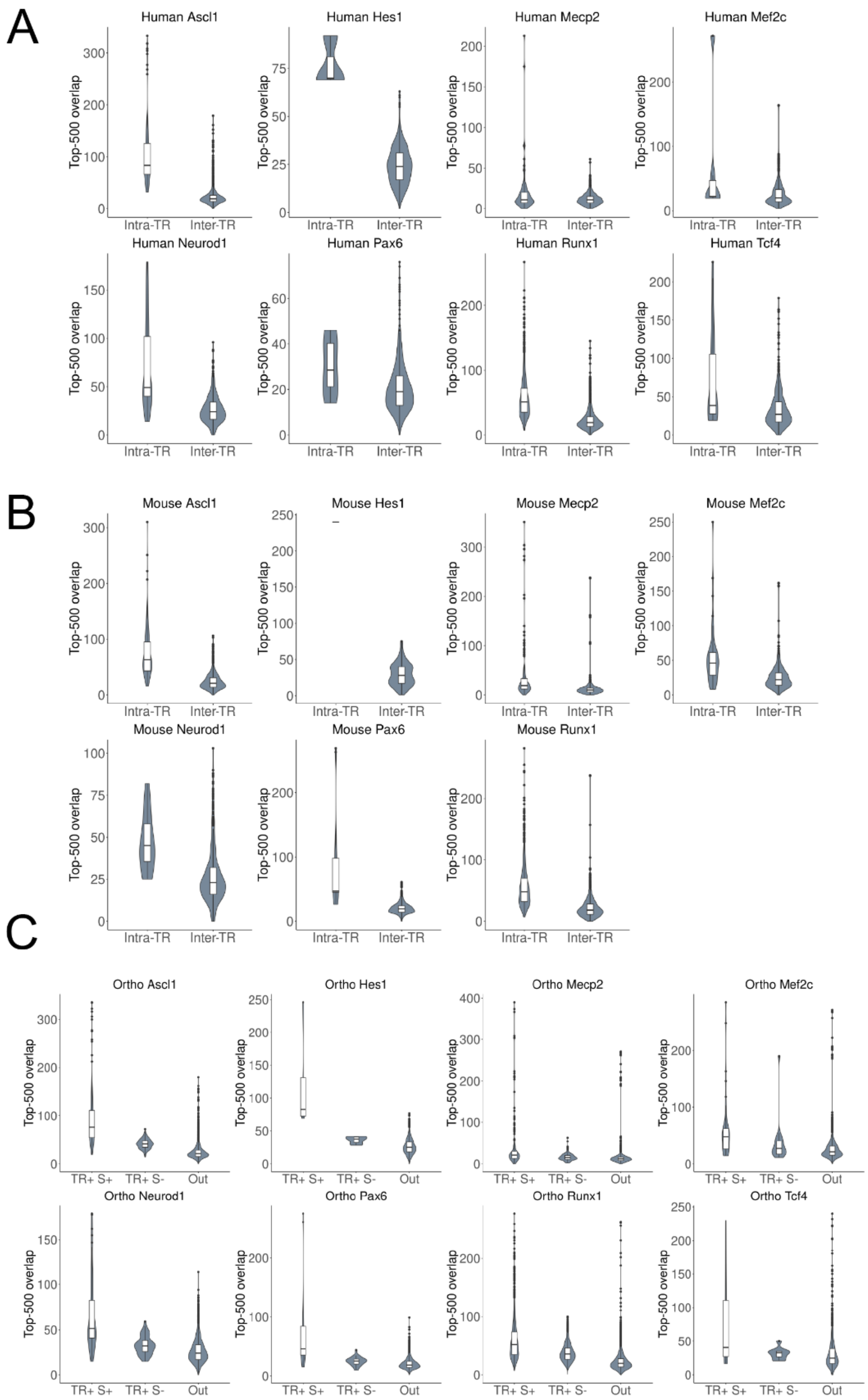


**Figure S2**. Distribution of the count of overlapping genes when selecting the top 500 genes by binding score when comparing ChIP-seq experiments targeting the same TR (intra-TR) or a different TR (inter-TR) for (A) human, (B) mouse, and (C) both species when using only high-confidence 1:1 orthologous genes. Abbreviations in C: TR+ (intra-TR), S+ (within-species), S- (cross-species), Out (collapsing inter-TR & intra-species with inter-TR & cross-species).

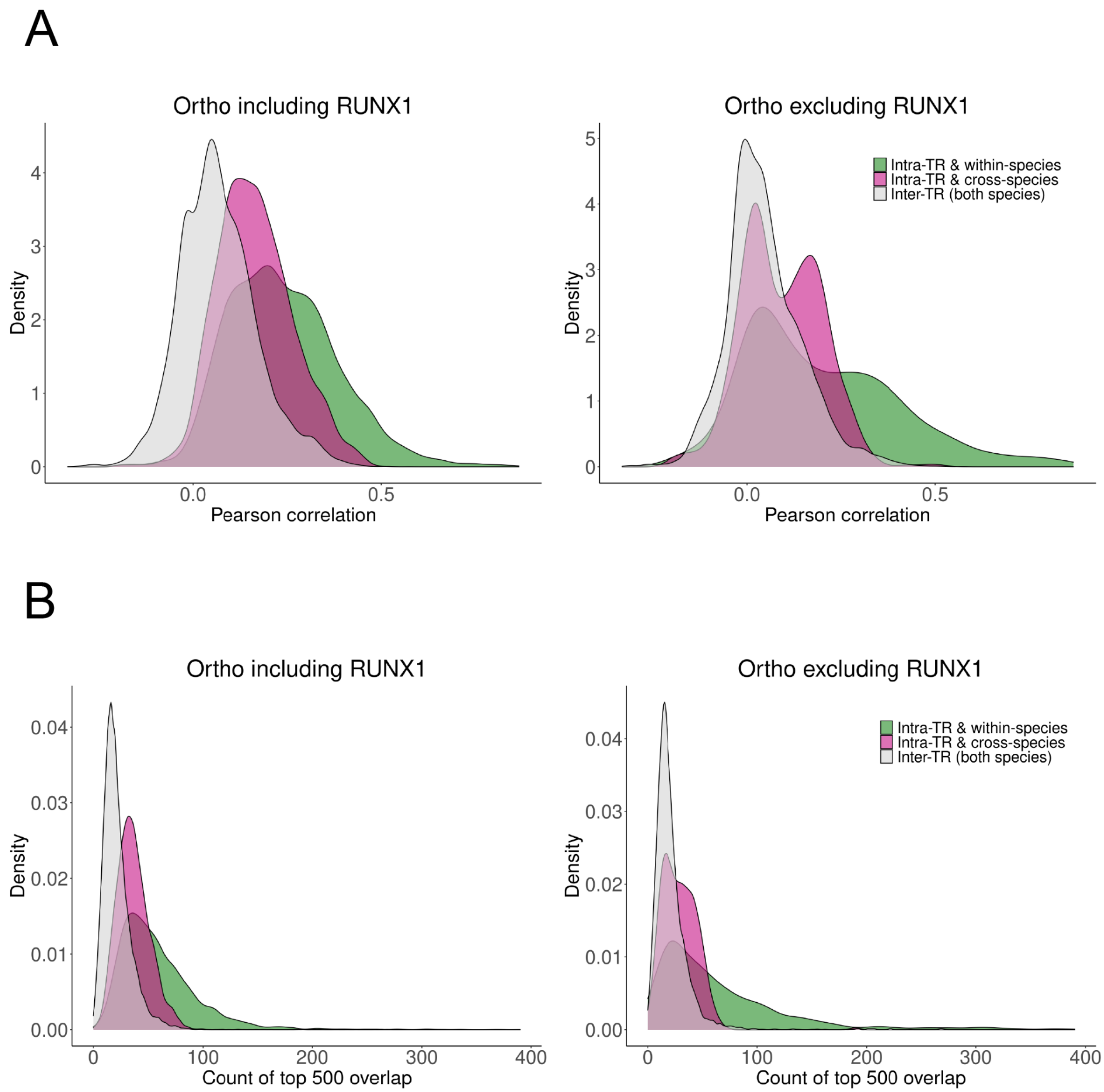


**Figure S3.** ChIP-seq experiment similarity of orthologs with and without RUNX1. (A) Left panel: As in Figure 2C, the distribution of binding score correlations between ChIP-seq experiments targeting the same TR (intra-TR) versus different TRs (inter-TRs), considering only orthologous genes. Right panel: The same comparison, except excluding all RUNX1 experiments. (B) Same as in A, except showing the density plots of the count of top 500 overlapping genes by binding score between pairs of experiments.

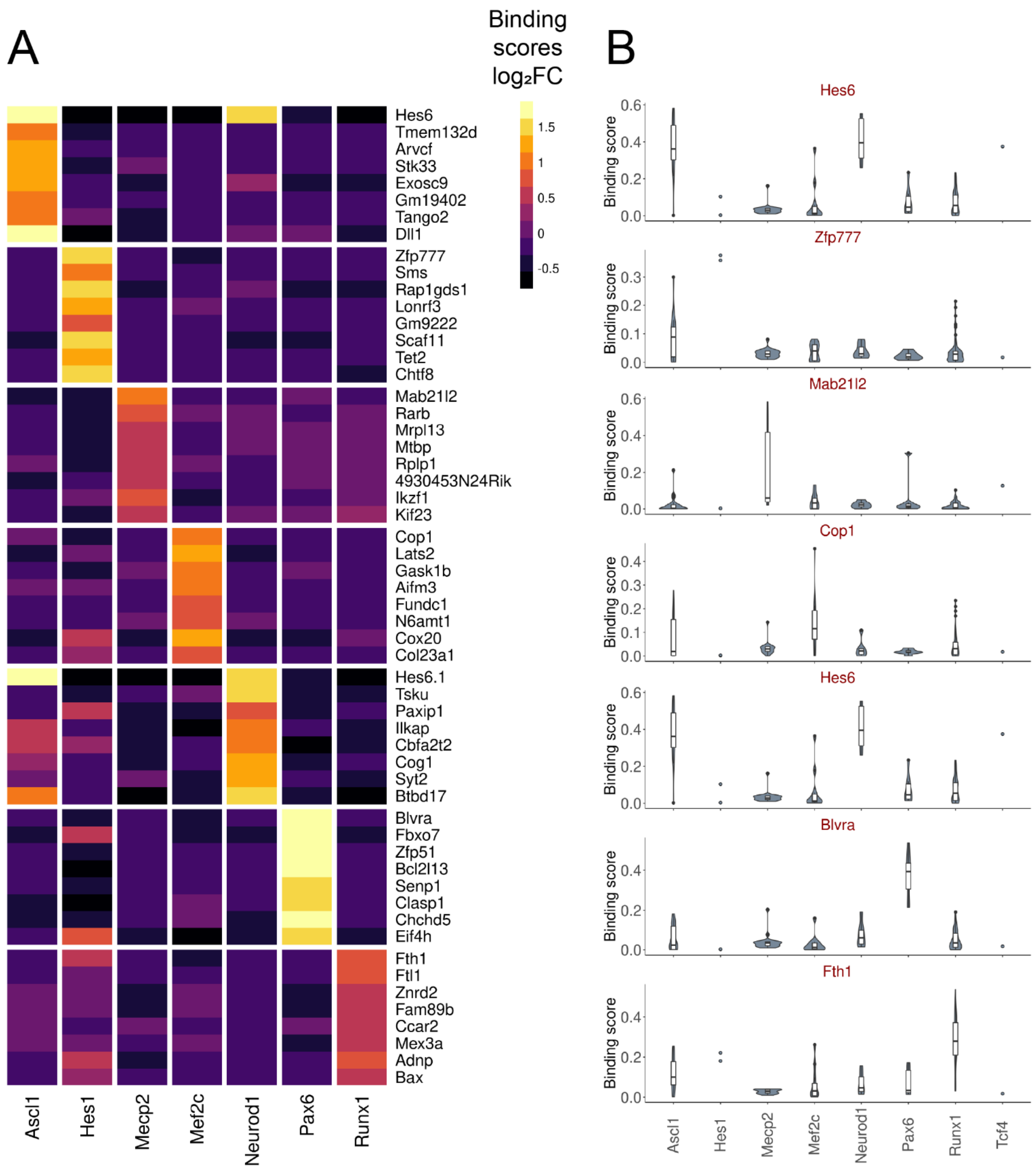


**Figure S4**. Specifically bound genes in mouse. (A) $\text{Log}_2$ fold changes of binding scores for the top eight ranked genes (by p-value) for each TR in mouse experiments using a mixed effect linear model. (B) Distribution of binding scores for the most significant gene for each TR extracted from the same model as in (A). Note that *Hes6* was the most significant gene in both the Ascl1 and the Neurod1 contrast.

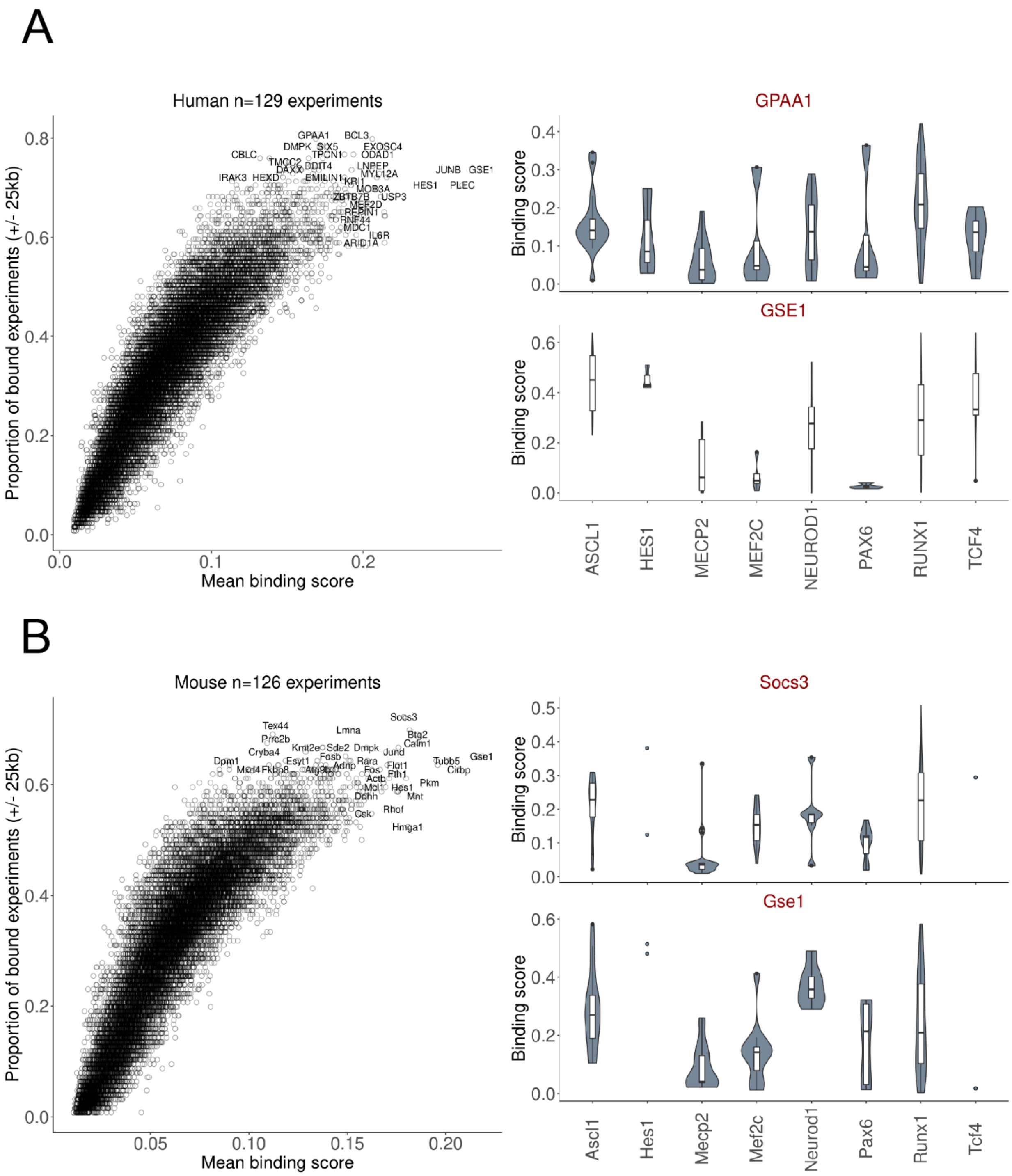


**Figure S5**. Demonstration of genes with high aggregated scores in the current ChIP-seq corpus. (A) Left panel scatter plot: every point is a gene, where the y-axis shows the proportion of human ChIP-seq experiments that had a peak within 25 kbp of the gene TSS (binary binding assignment), and the x-axis shows the mean of the continuous binding score across all experiments. Right panel: distribution of binding scores for the top gene by proportion bound (GSAA1) and by mean binding score (GSE1). (B) Same as in (A), but for mouse experiments.

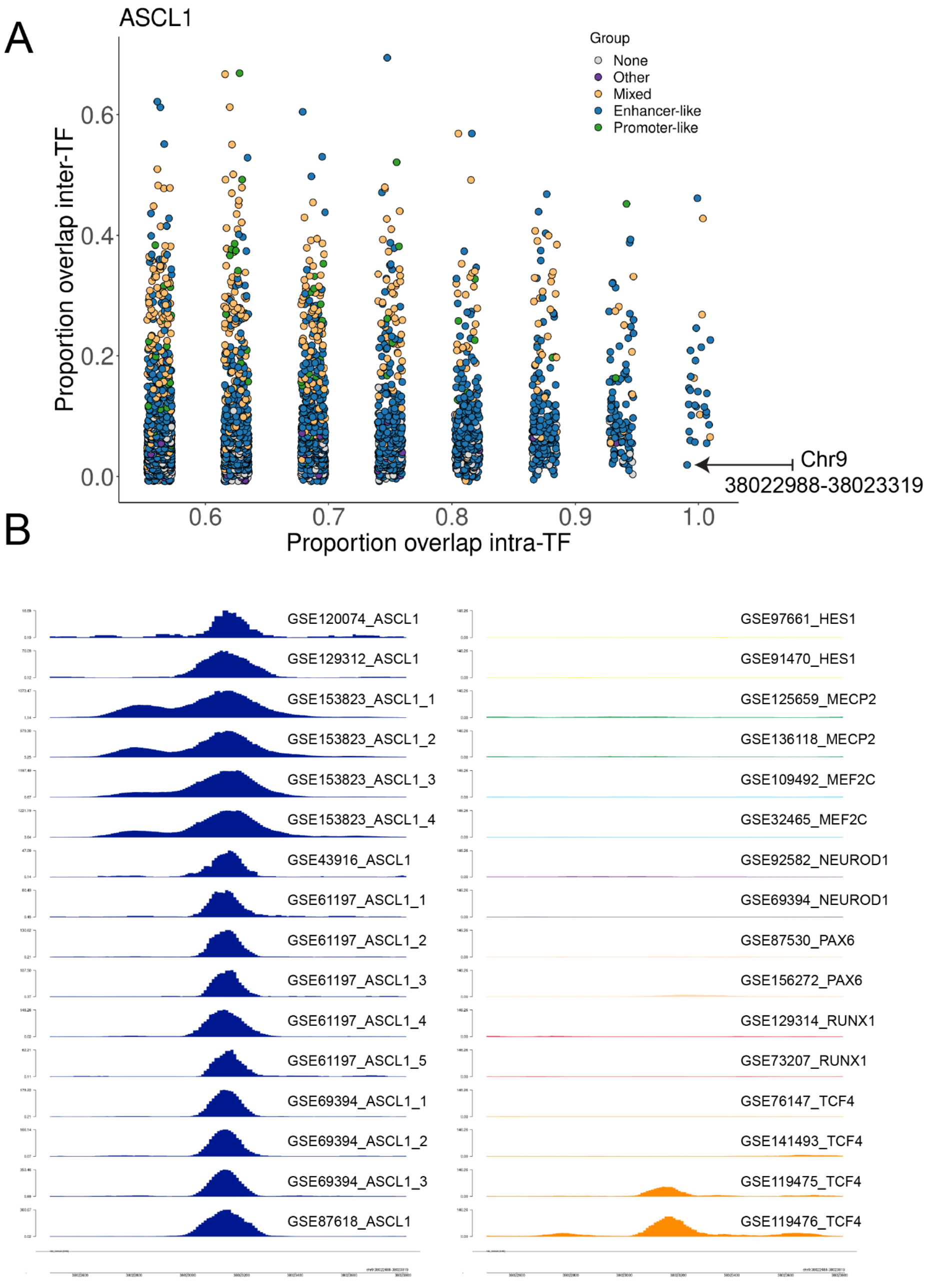


**Figure S6.** ASCL1 frequently bound loci. (A) Each point represents a human ASCL1-bound region, restricted to regions bound in at least half of ASCL1 experiments. The x-axis is the proportion of human ASCL1 experiments (n=16) in which that region is bound, and the y-axis is this proportion for non-ASCL1 experiments (n=113). Points are colored by ENCODE cCRE definitions, binning each enhancer or promoter subgroup (seen in legend of Figure 3A), and designating non-ELS/PLS elements as 'Other.' Mixed refers to regions straddling two cCRE groups. (B) Trackplot highlighting an intronic region of *SHB*, indicated with an arrow in (A), that had a peak in all ASCL1 experiments and only in 2/113 non-ASCL1 experiments. Left panel shows all ASCL1 experiments, right panel shows two sampled experiments for the other TRs, as well as the two TCF4 experiments in which this region had a peak called. Note that the left panel has variable y-axes limits (reflecting read enrichment) to fit all the peaks, while the right panel y-axes limits are fixed to the median of the maximum ASCL1 intensities to demonstrate the relative lack of signal and to fit the two TCF4 peaks.

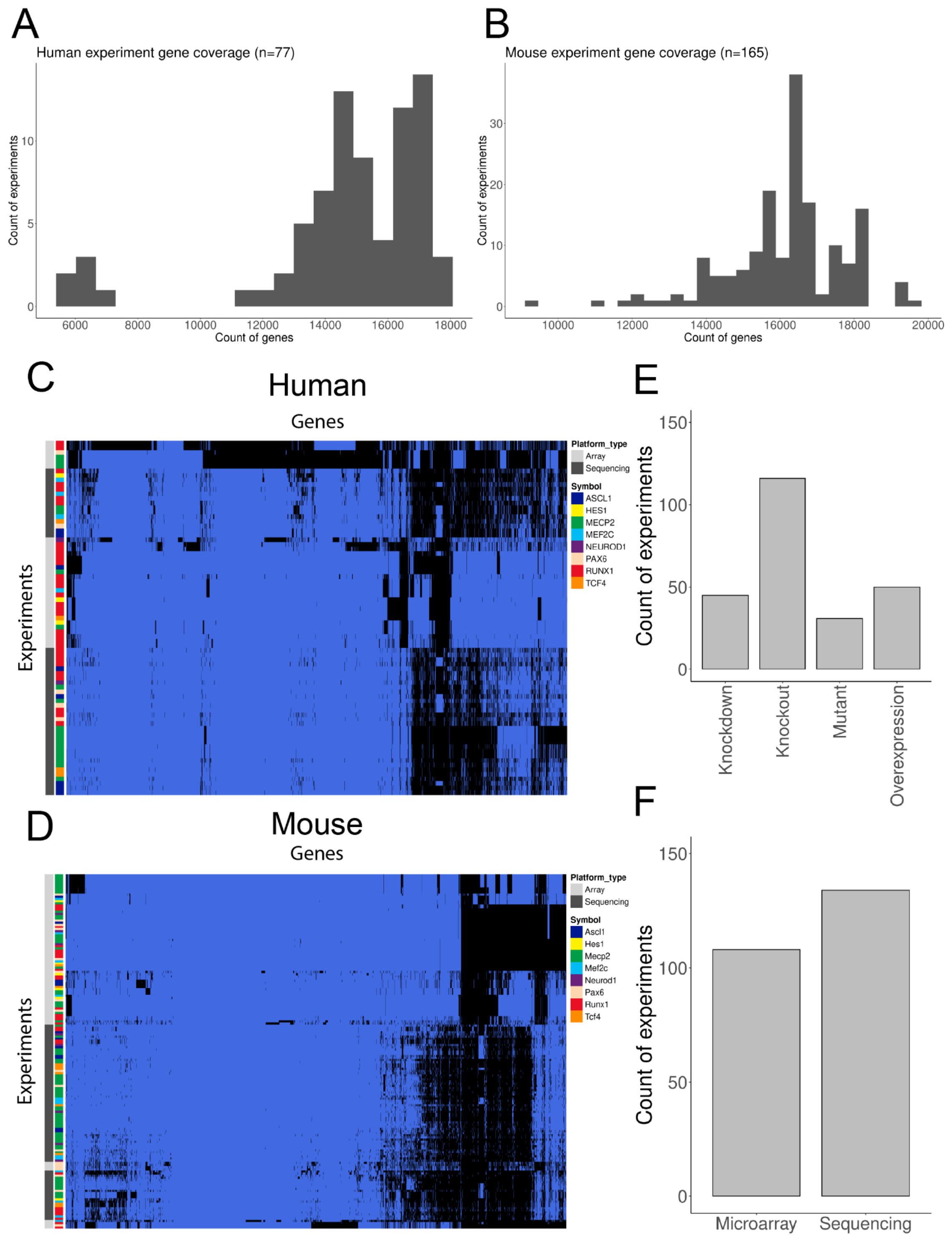


**Figure S7.** TR perturbation experiment overview. (A-B) Histograms of the count of genes measured in human and mouse perturbation experiments. For microarray platforms, genes that were not measured are not on the platform or were filtered in QC. For RNA-sequencing, genes that were not measured were not appreciably detected in the tested context. (C-D) Binary heatmap of genes measured across human (C) and mouse (D) experiments; blue = measured, black = not measured. (E-F) Barcharts for the count of experiments by perturbation strategy (E) or expression technology (F).

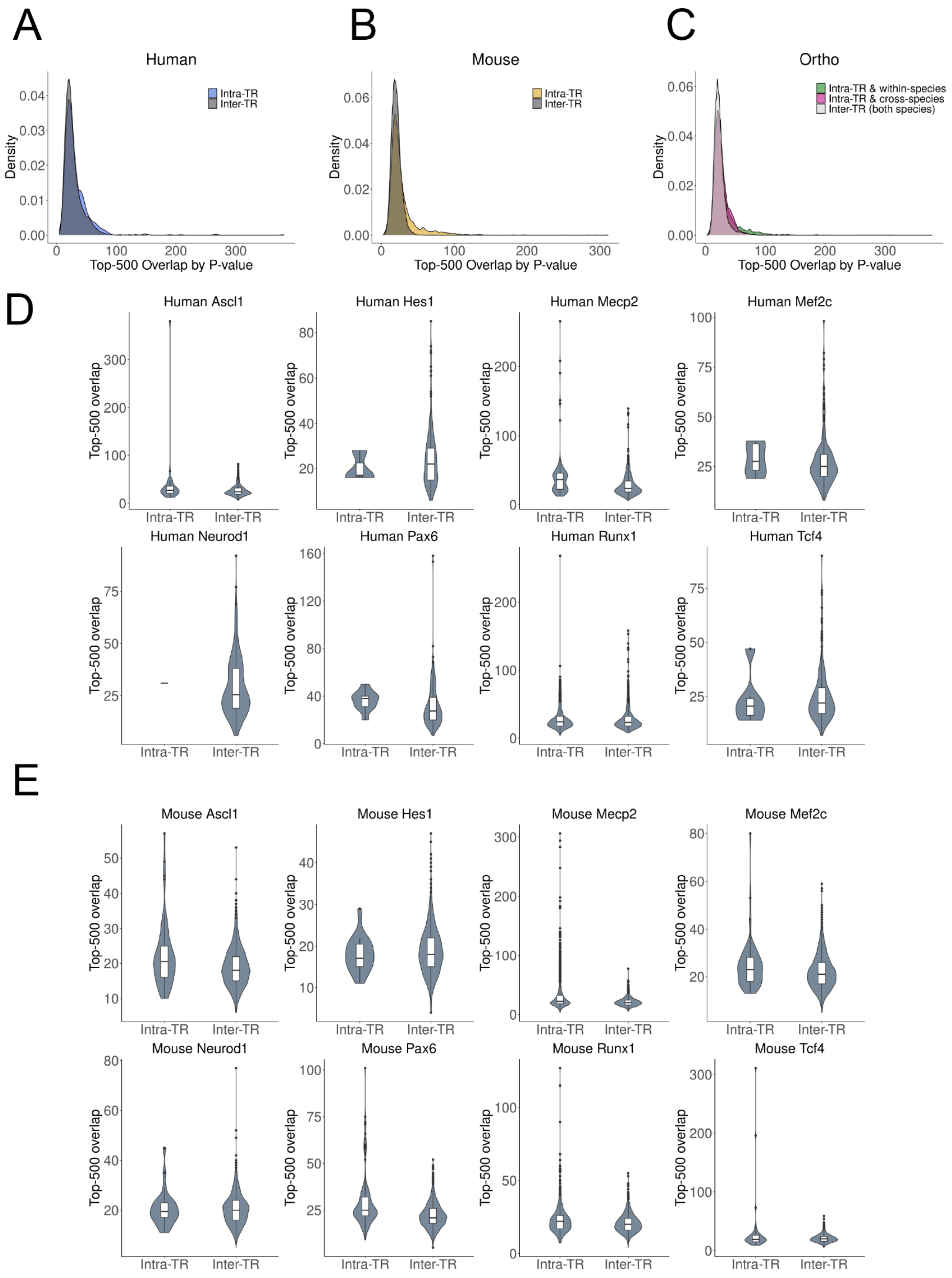


**Figure S8.** TR perturbation experiment similarity. (A-C) Density plots of the count of overlapping genes between perturbation experiments, selecting for the top 500 mutually-measured genes sorted by p-value from the differential expression analysis. (D) Distribution of the counts from (A) broken down by TR. (E) Distribution of the counts from (B) broken down by TR.

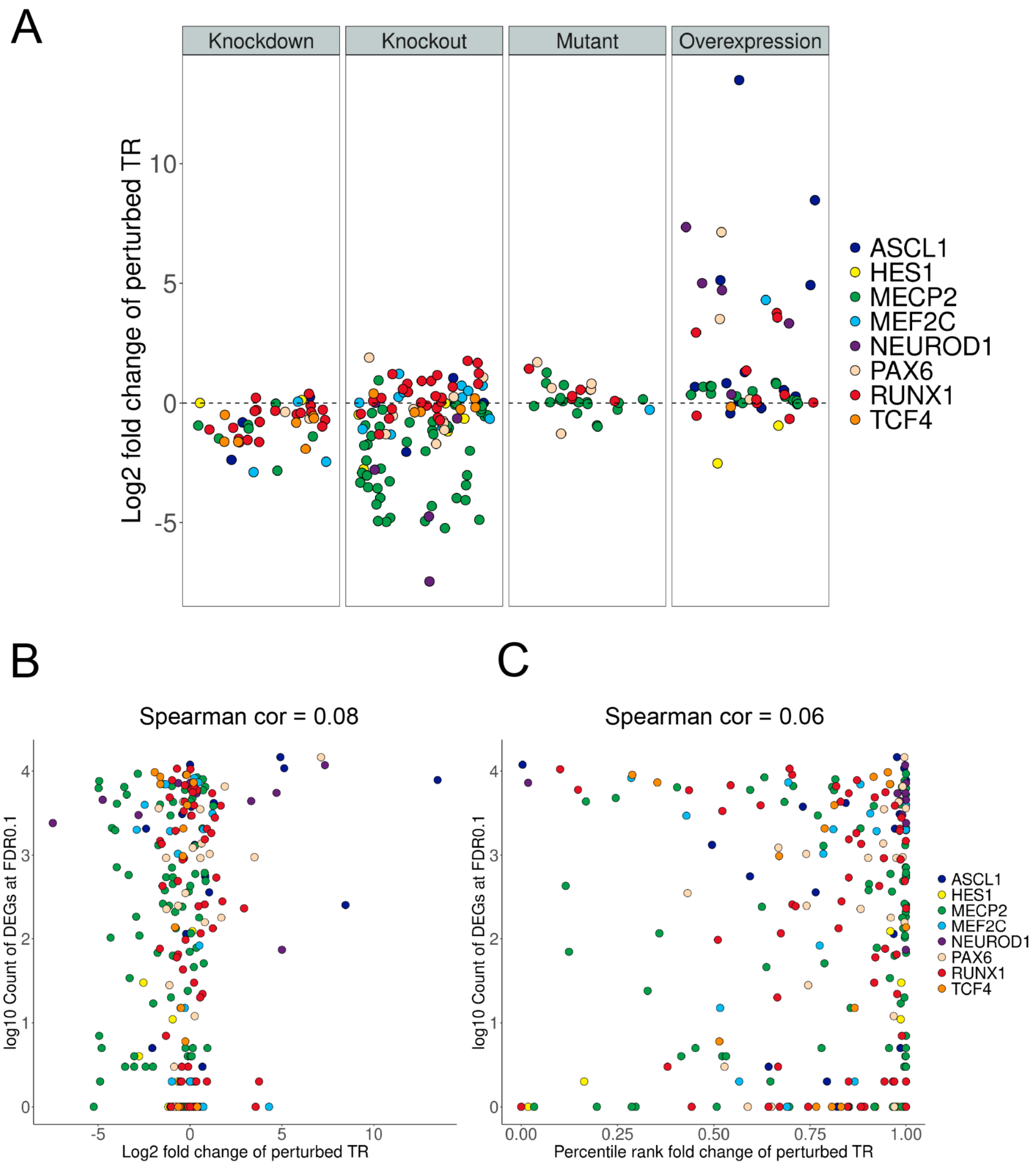


**Figure S9.** Perturbation effect sizes. (A) Each point represents the $\log_2$ fold change of the perturbed TR in mouse and human perturbation experiments, coloured by TR and grouped by perturbation strategy. (B-C) Relationship between the effect size of the perturbed TR and the count of differentially expressed genes in that experiment. (B) The fold change of the perturbed TR is shown, while (C) shows the percentile rank of the absolute fold change of the perturbed TR (where 1.0 = the gene with the highest absolute fold change in that experiment).

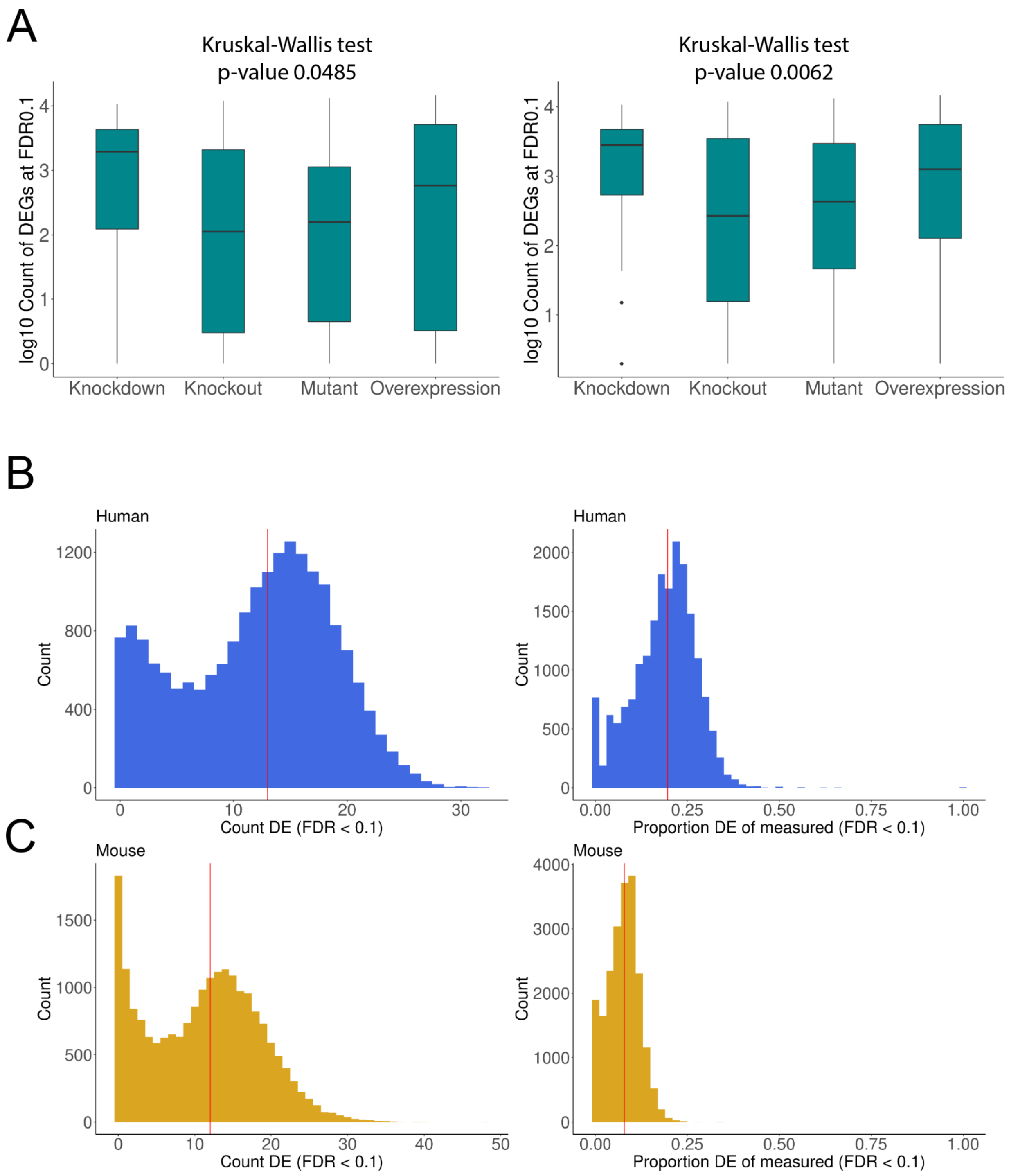


**Figure S10.** Count of differentially expressed genes (DEG). (A) Count of DEGs by perturbation strategy, either including experiments with no DEGs (left panel) or excluding experiments with no DEGs (right panel). (B) Left panel: histogram of the count of times a gene was measured as DE across human experiments (n=77); Right panel: instead of raw counts, the histogram shows the proportion of DE measurements relative to the number of experiments in which the gene was measured. (C) Same as in (B), except for mouse experiments (n=165).

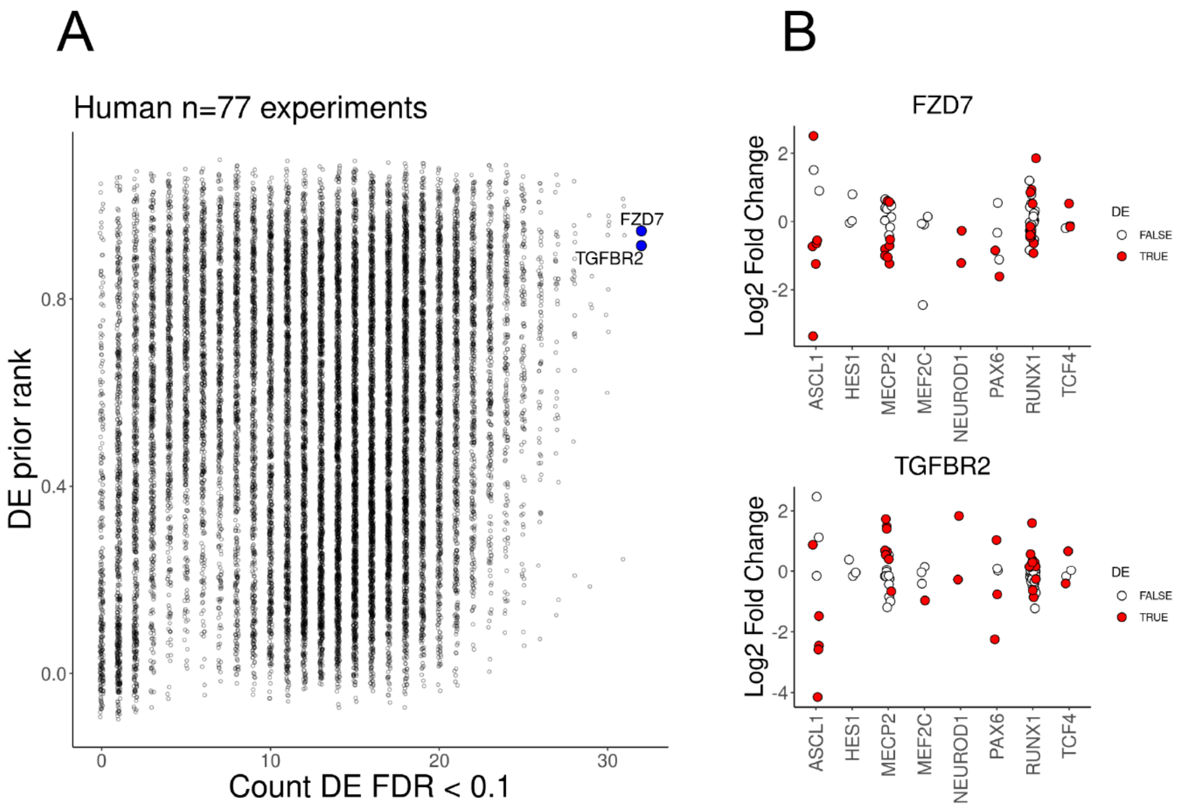


**Figure S11.** Frequently DE genes. (A) Relationship between the count of times a gene was DE across all human experiments for the eight TRs, and the DE prior ranking, as in Figure 5A. Highlighted are the two genes with the highest DE counts. (B) FC distributions for the genes highlighted in (A), coloured by DE status.

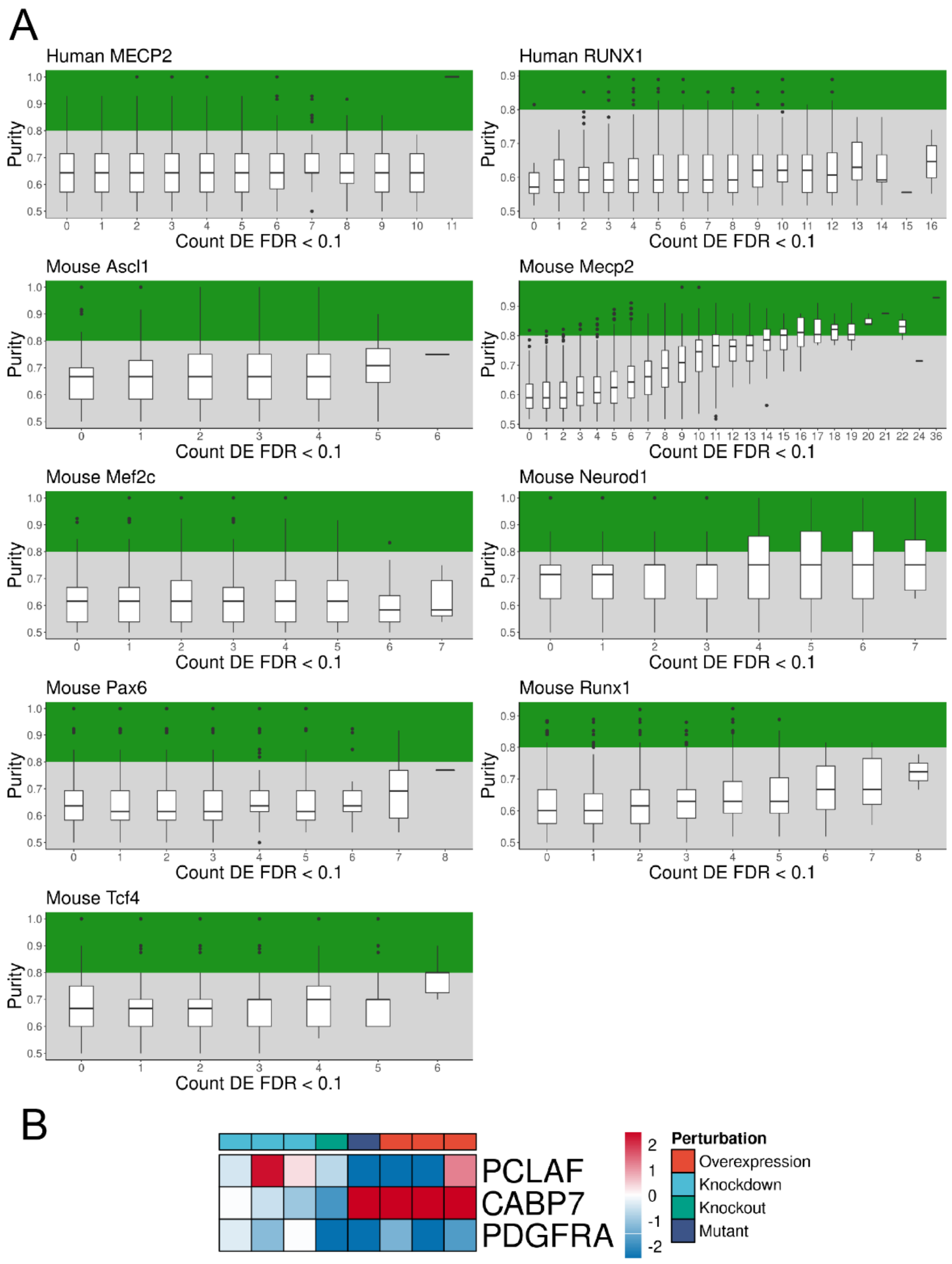


**Figure S12.** Genes typically show variable changes in FC direction across TR perturbation experiments. *Purity* is a measurement of the consistency of a gene's FC direction within TR gain/loss of function experiments. (A) Demonstrates the TR-specific distributions of gene *Purity* scores, stratified by how often the genes were measured as DE across each set of TR perturbation experiments. Only TRs with at least eight non-mutant experiments were kept to alleviate issues of sample size. Genes with consistent FC direction were arbitrarily defined as having *Purity* > 0.8 (green background). (B) Shows three genes with strong human ASCL1 DE evidence (DE > 4/8 ASCL1 experiments), as in Figure 5D. *PCLAF* has low *Purity* (0.57) due to variation in its FC direction. *CABP7* has perfect *Purity* and *Signed purity* (1.00), while *PDGFRA* has a *Signed purity* of -1.00 as the FC direction is the same for gain/loss of function experiments.

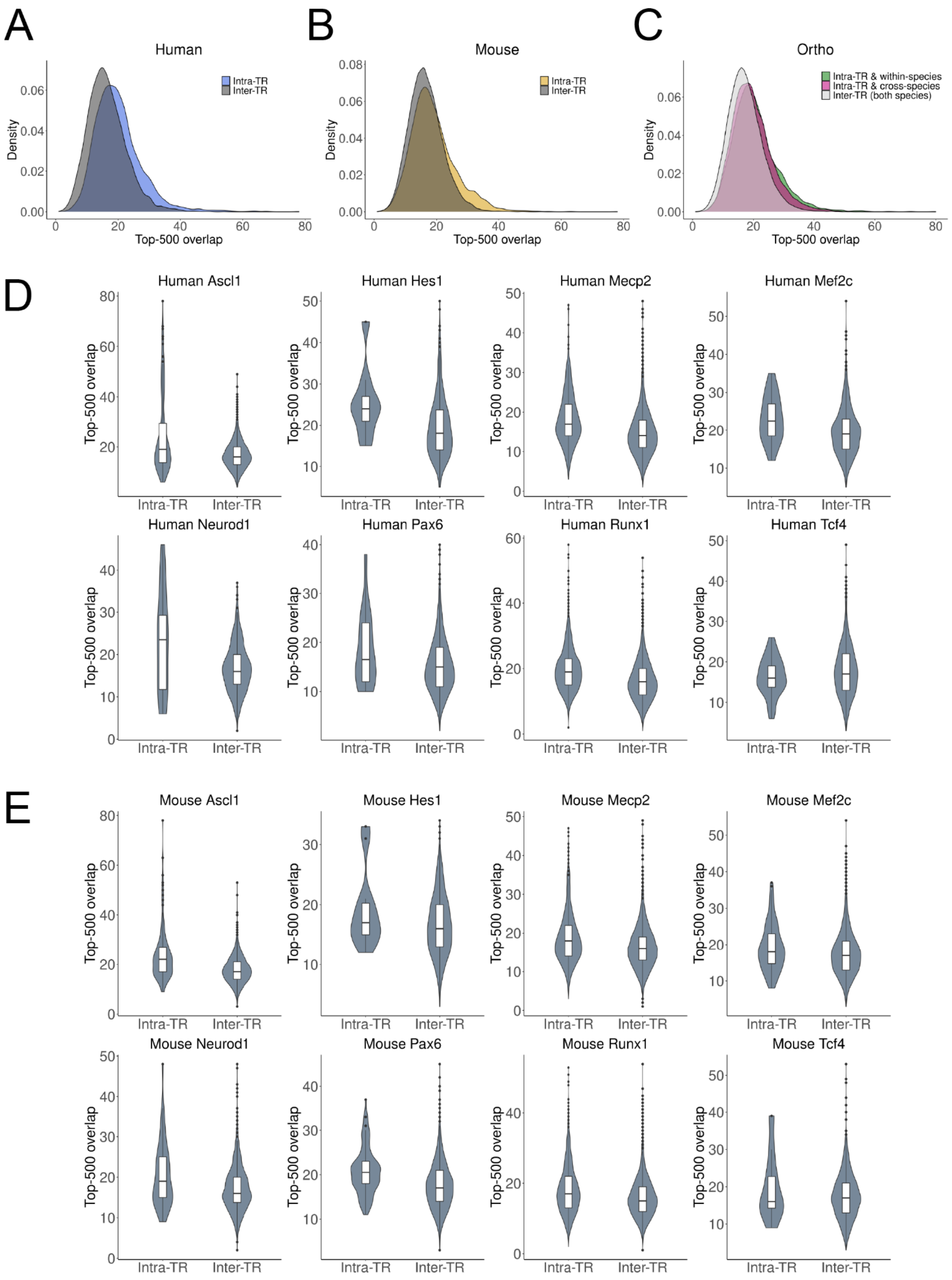


**Figure S13.** Top scoring gene overlap between ChIP-seq and TR perturbation experiments. (A-C) Density plots of the count of intersecting genes between ChIP-seq and perturbation experiments, selecting for the top 500 genes by binding score (ChIP-seq) or by p-value from the DE analysis (perturbation). (D) Distribution of the counts from (A) broken down by TR. (E) Distribution of the counts from (B) broken down by TR.

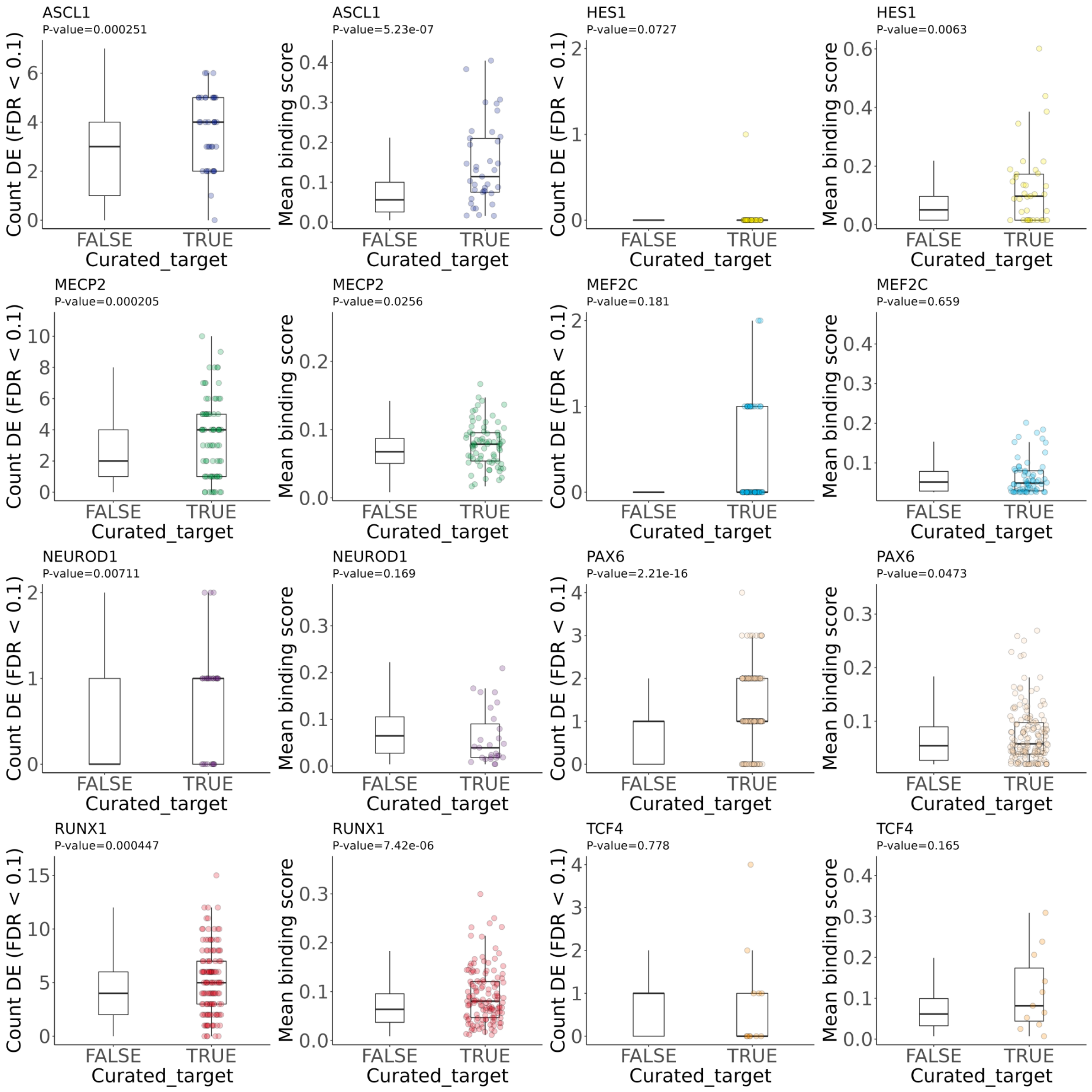


**Figure S14.** Human aggregate scores by curation status. Boxplots show the aggregated genomic experiment gene scores for gene status in the low-throughput evidence resource. P-values are from the Wilcoxon rank sum test.

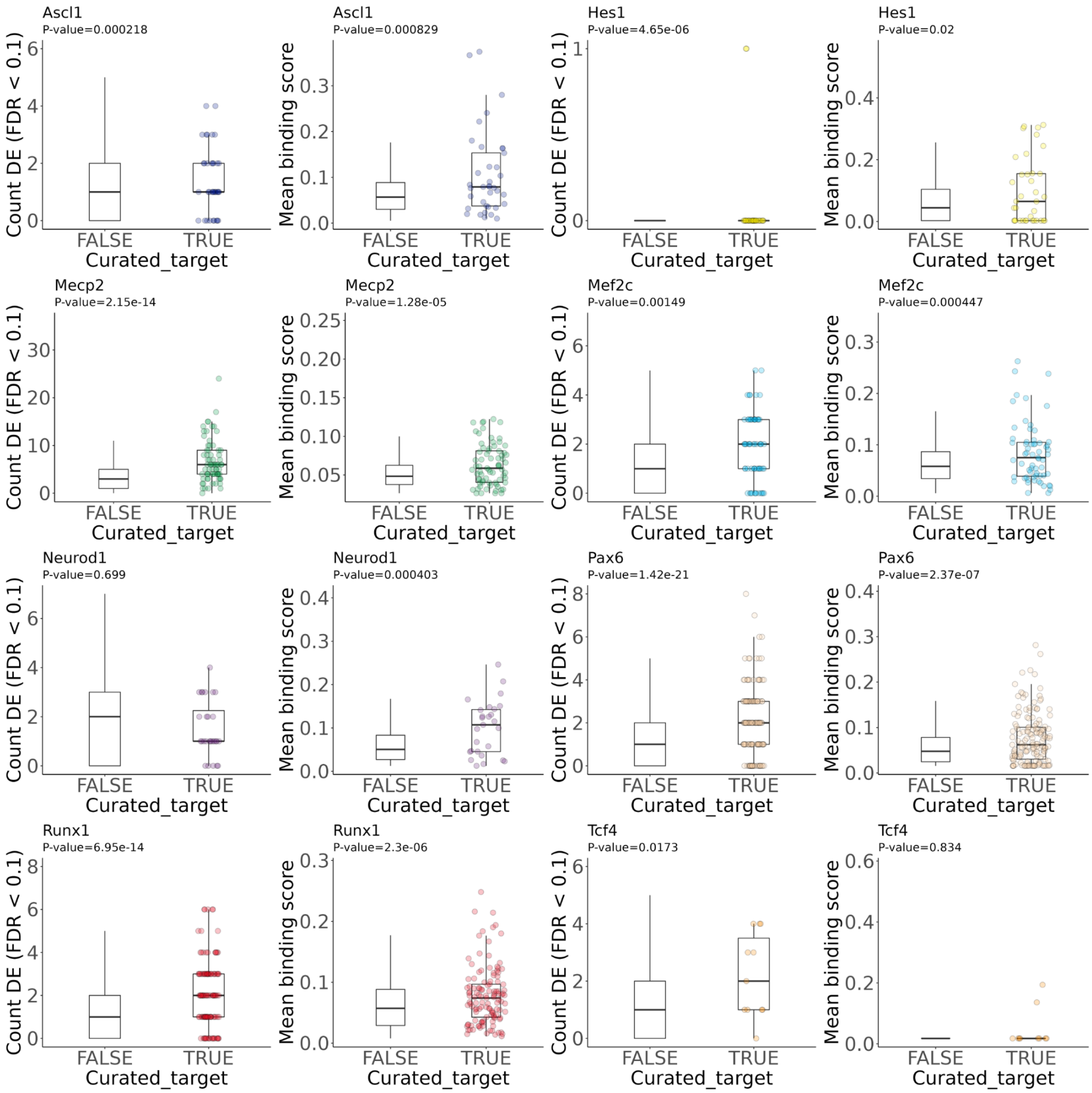


**Figure S15.** Mouse aggregate scores by curation status. Boxplots show the aggregated genomic experiment gene scores for gene status in the low-throughput evidence resource. P-values are from the Wilcoxon rank sum test.

A

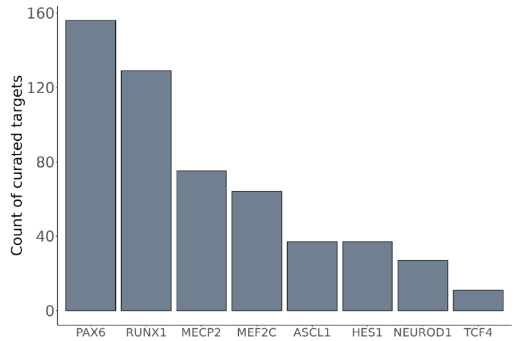


B

**AUPRC — Human**

| TR | Integrated | Binding | Perturbation |
| --- | --- | --- | --- |
| ASCL1 | 0.053 | 0.041 | 0.019 |
| HES1 | 0.050 | 0.060 | 0.020 |
| MECP2 | 0.017 | 0.013 | 0.019 |
| MEF2C | 0.017 | 0.012 | 0.028 |
| NEUROD1 | 0.021 | 0.020 | 0.022 |
| PAX6 | 0.028 | 0.014 | 0.036 |
| RUNX1 | 0.026 | 0.019 | 0.018 |
| TCF4 | 0.074 | 0.051 | 0.138 |

**Sampled AUPRC greater than observed — Human**

| TR | Integrated | Binding | Perturbation |
| --- | --- | --- | --- |
| ASCL1 | 0.000 | 0.003 | 0.084 |
| HES1 | 0.000 | 0.000 | 0.048 |
| MECP2 | 0.014 | 0.058 | 0.010 |
| MEF2C | 0.033 | 0.864 | 0.011 |
| NEUROD1 | 0.743 | 0.962 | 0.312 |
| PAX6 | 0.000 | 0.382 | 0.000 |
| RUNX1 | 0.000 | 0.018 | 0.018 |
| TCF4 | 0.001 | 0.186 | 0.000 |

**Percentile AUPRC relative to individual experiments — Human**

| TR | Integrated | Binding | Perturbation |
| --- | --- | --- | --- |
| ASCL1 | 1.000 | 1.000 | 0.316 |
| HES1 | 1.000 | 1.000 | 0.333 |
| MECP2 | 0.950 | 0.716 | 0.993 |
| MEF2C | 0.750 | 0.333 | 0.917 |
| NEUROD1 | 0.875 | 0.344 | 0.938 |
| PAX6 | 1.000 | 0.310 | 1.000 |
| RUNX1 | 0.981 | 0.883 | 0.860 |
| TCF4 | 0.918 | 0.510 | 1.000 |

**AUPRC — Mouse**

| TR | Integrated | Binding | Perturbation |
| --- | --- | --- | --- |
| Ascl1 | 0.051 | 0.056 | 0.022 |
| Hes1 | 0.052 | 0.019 | 0.066 |
| Mecp2 | 0.027 | 0.015 | 0.033 |
| Mef2c | 0.017 | 0.017 | 0.013 |
| Neurod1 | 0.021 | 0.023 | 0.019 |
| Pax6 | 0.043 | 0.017 | 0.038 |
| Runx1 | 0.030 | 0.015 | 0.021 |
| Tcf4 | 0.047 | 0.046 | 0.048 |

**Sampled AUPRC greater than observed — Mouse**

| TR | Integrated | Binding | Perturbation |
| --- | --- | --- | --- |
| Ascl1 | 0.000 | 0.000 | 0.034 |
| Hes1 | 0.000 | 0.058 | 0.000 |
| Mecp2 | 0.007 | 0.040 | 0.000 |
| Mef2c | 0.022 | 0.021 | 0.289 |
| Neurod1 | 0.066 | 0.024 | 0.939 |
| Pax6 | 0.000 | 0.074 | 0.000 |
| Runx1 | 0.001 | 0.096 | 0.018 |
| Tcf4 | 0.194 | 0.613 | 0.062 |

**Percentile AUPRC relative to individual experiments — Mouse**

| TR | Integrated | Binding | Perturbation |
| --- | --- | --- | --- |
| Ascl1 | 0.910 | 0.927 | 0.515 |
| Hes1 | 0.950 | 0.250 | 1.000 |
| Mecp2 | 0.969 | 0.615 | 0.998 |
| Mef2c | 0.832 | 0.832 | 0.467 |
| Neurod1 | 0.452 | 0.548 | 0.000 |
| Pax6 | 0.934 | 0.331 | 0.887 |
| Runx1 | 0.981 | 0.548 | 0.859 |
| Tcf4 | 0.000 | 0.000 | 0.000 |

**Figure S16.** Detailed evaluation metrics for TR target prediction. (A) Count of unique targets in the low-throughput target resource. For evaluation we accepted all low-throughput modes of evidence and included orthologous gene targets conducted in either species. (B) Left panel: AUPRC values are as demonstrated in Figure 6A,B. Mid panel: The proportion of sampled targets whose AUPRC exceeded the observed aggregated AUPRC, as in Figure 6C. 1000 samples size matched for each TR were drawn from the entire curated resource and evaluated with the integrated ranking. Right panel: The percentile of the aggregated AUPRCs relative to the distribution of AUPRCs generated by treating each contributing experiment as its own ranking, as in Figure 6D.

#### Low-throughput curated target resources

Our evaluation of the genomic rankings was facilitated by resources that curated biochemical assays probing TR-target interactions. These include low-throughput perturbation assays (such as qPCR), binding assays (ChIP, EMSA), and reporter assays (such as luciferase reporters). The majority of the curated interactions (60%) are uniquely identified from our previous work, Chu et al., 2021 (here, Chu2021), along with additional interactions that have been curated since original publication. These interactions are also noted as originating from Chu2021, although a manuscript of the increased coverage is in preparation. Additionally, we include targets that have been curated by other resources, which was organized in Chu2021. TRRUST (Han et al., 2018) was by far the most prevalent of these resources. Interactions curated by Chu2021 and/or TRRUST comprise 95% of the low-throughput TR-target relationships in this study. In decreasing order of prevalence, the remaining resources that were included are TFe (Yusuf et al., 2012), ENdb (Bai et al., 2020), CytReg (Carrasco Pro et al., 2018), HTRIdb (Bovolenta et al., 2012), InnateDB (Lynn et al., 2008), TFactS (Essaghir et al., 2010), and ORegAnno (Lesurf et al., 2016).

Finally, we note that multiple of the TCF4 interactions found when combining all of the curated resources were actually TCF7L2, which is frequently referred to as TCF4 (T-Cell Factor 4) in the literature. We purged all such examples, resulting in a dramatic decrease in the number of TCF4 curated targets.

### Overview of TR-targets

RUNX1 was the best represented TR overall, with many of the data sets from this collection coming from either blood progenitors or blood cancer cell lines. Correspondingly, we found examples of hematopoietic genes with curated evidence and a high integrated ranking, such as the erythroid TF *NFE2* (human: 94th, mouse: 13th; Wang et al., 2010) and immune adhesion subunit *ITGB2* (human: 145th, mouse: 20th; Puig-Kröger et al., 2003). We also found many genes involved in cellular motility and cytoskeletal functionality among the top ranked. Some, like myosin heavy chain 9 (*MYH9;* human: 229th, mouse: 219th), had curated evidence (Bluteau et al., 2012). Most however did not, such as plectin (*PLEC*; human: 2nd, mouse: 17th), noted to be frequently implicated in cancer (Perez et al., 2021*)*, as well as lamin A/C (*LMNA*; human: 1,628th, mouse: 27th), vimentin (*VIM;* human: 5th, mouse: 49th), and paxillin (*PXN;* human: 93rd, mouse: 292nd).

We caution that the aggregated rankings did not always prioritize curated/known interactions, such as the important hematopoietic TF *SPI1* (human: 2,342nd, mouse: 1,477th; Huang et al., 2008). *SPI1* had moderately elevated RUNX1 binding in both species, but weaker perturbation evidence lowered its integrated rankings. We also note that RUNX1 was ranked first as its own target in both species. While autoregulatory functionality has been shown for this TR (Nottingham et al., 2012), the rankings will prioritize each respective TR by virtue of these genes being frequently DE in their perturbation experiments. Nevertheless, the rankings also highlighted orthologous genes that had strong evidence in both species but are rarely studied, such as *MOB3A (*human: 10th, mouse: 101st), believed to be involved in the angiogenic Hippo pathway-YAP/TAZ (Boopathy and Hong 2019; Dutchak et al., 2022). Finally, we highlight that *PHF19* (human: 7th) was recently identified in a crowd-sourced competition as an aggressive marker of multiple myeloma (Mason et al., 2020), suggesting further potential oncogenic activity of RUNX1 misregulation.

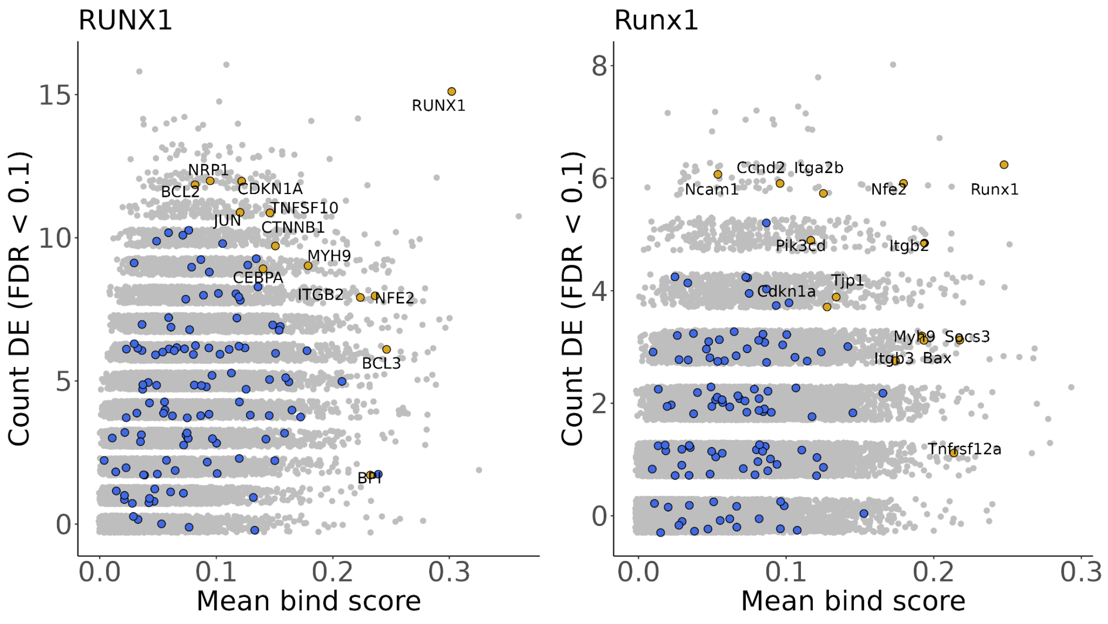

MECP2 was the next most represented TR and had the most brain-relevant experiments. While the TSS-based logic of the binding score metric is likely better suited to the TFs, the rankings still provided relevant examples. *Sdk1* (human: 1,630th, mouse: 41st) and *Auts2* (human: 6,236th, mouse: 10th) both ranked highly in mouse. A recent murine Mecp2 study demonstrated that these genes are in a contacting chromosomal domain enriched for methylated cytosines (Clemens et al., 2020), adding a further line of evidence beyond the scope of this study. *Sdk1* has been studied for its role in retinal development and cocaine addiction behavior (Scobie et al., 2014; Yamagata and Sanes 2019), while the connection between *Mecp2* and *Auts2* in particular has garnered interest for its involvement in neurodevelopmental and psychiatric outcomes (Pang et al., 2021).

The top curated target by human rankings was *BDNF* (human: 29th, mouse: 2,968th; Abuhatzira et al., 2007), although this was driven almost exclusively by the perturbation evidence. In mouse it was *Aff1* (human: 16,335th, mouse: 30th; Urdinguio et al., 2008), which was well supported by both data types. We also note that the curated target *Irak1* (Urdinguio et al., 2008), which neighbours the *Mecp2* loci, had excellent perturbation evidence in mouse but essentially no binding evidence in our framework. It has been posited that *Mecp2* regulates *Irak1* through both indirect (Urdinguio et al., 2010) and direct (Kishi et al., 2016) means. Examples of candidates without curated evidence include the rarely studied *CDS1* (human: 59th, mouse: 56th), which is involved in lipid droplet growth (D'Souza et al., 2014; Xu et al., 2019). Interestingly, aberrant lipid growth has been documented in murine Rett syndrome models (Vashi et al., 2021). Other examples of high ranking targets relevant to brain health but lacking curated evidence include *SHANK2* (human: 621st, mouse: 193rd; Zaslavsky et al., 2020), *PLXNA2* (human: 60th, mouse: 288th; Kong et al., 2016; Pijuan et al., 2021), *CNTNAP2* (human: 1,326th, mouse: 111th; Xing et al., 2019), *TENM2* (human: 3,581st, mouse: 219th; Wray et al., 2018), and *TENM3* (human: 5,991st, mouse: 15th; Singh et al., 2019).

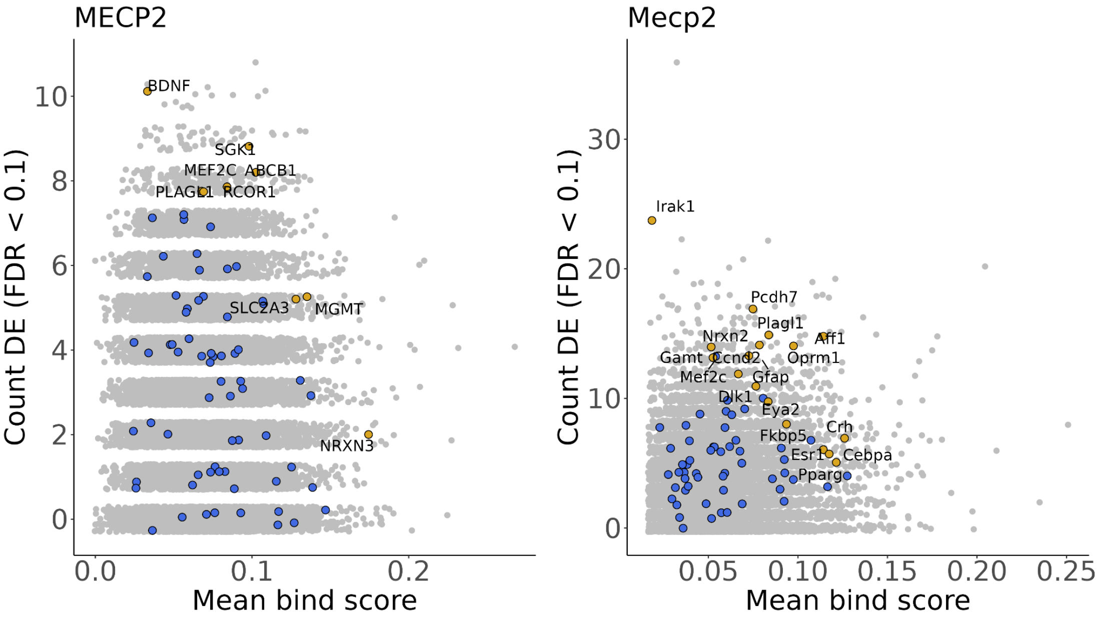

ASCL1 experiments can be broadly characterized as deriving from embryonic systems (some from primary tissues) in mouse, while human experiments were overwhelmingly carried out in cancer cell lines. Highly ranked and previously described targets include Notch pathway effectors *DLL1* (human: 2nd, mouse: 5th), *DLL3* (human: 70th, mouse: 8th), *DLL4* (human: 35th, mouse: 1,168th), *HES6* (human: 57th, mouse: 3rd) and *JAG2* (human: 5th, mouse: 4,595th) (Castro et al., 2006; Henke et al., 2009; Johansson et al., 2009; Nelson et al., 2009; Castro et al., 2011; Augustyn et al., 2014; Misra et al., 2014). Notch effector *LFNG* was also highly ranked (human: 9th, mouse: 138th) but was not in the curated resources. Similarly, the cell-cycling *CDC25B* had robust evidence in both data types for both species (human: 3rd, mouse: 1st); this gene was not in the curated resource but previously had been noted as a candidate target (Castro et al., 2006; Castro et al., 2011).

Other examples of candidates lacking curated evidence but have been putatively linked to ASCL1 include the repressive TFs *ID1* (human: 11th, mouse: 27th) and *ID3* (human: 12th, mouse: 13th) (Jorstad et al., 2020), as well as *ZBTB18* (human: 17th, mouse: 29th) (Wapinski et al., 2013). Our work supports previous connections made between ASCL1 and the transcription repressor *CBFA2T3* (human: 442nd, mouse: 2nd) (Alishahi et al., 2009; Aaker et al., 2010), and novelly links ASCL1 to the epilepsy-implicated potassium channel gene *KCNH2* (human: 6th, mouse: 105th) (Bagnall et al., 2016 & 2017).

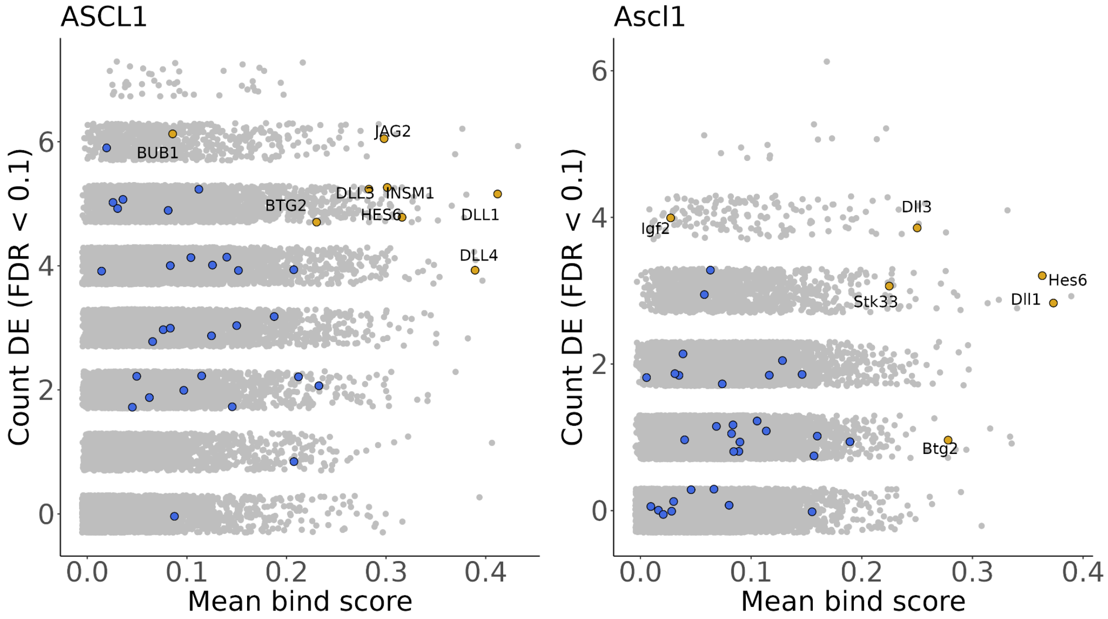

NEUROD1 experiments were typically conducted in embryonic stem cells, pancreatic tissues, or cancer cell lines. This TR had the second fewest targets in the curated resource, and only a handful of these also had high integrated ranks in one species alone, such as *Insm1* (human: 5,451st, mouse: 71st; Breslin et al., 2003). This gene is active in both neural and endocrine systems, and we found other examples of candidate targets that implicate NEUROD1's multi-tissue activity. *SRRM4* (human: 129th, mouse: 28th) encodes a crucial neurodevelopmental splicing factor recently implicated as contributing to dysregulated splicing in islets cells in a murine diabetic model (Quesnel-Vallières et al., 2015; Wilhelmi et al., 2021). Congenital disorders in Golgi functionality gene *COG1* (human: 296th, mouse: 44th) are associated with hypoglycemia and neurological impairment (Climer et al., 2018; Zhang et al., 2018; Huang et al., 2021). *CXXC4* (human: 5,465th, mouse: 3rd) has evidence supporting roles in beta cell differentiation and neuronal stem cell proliferation (Ahn et al., 2004; Guan et al., 2020), while *STXBP1* (human: 1,408th, mouse: 110th) has been linked to encephalopathies and enteroendocrine functionality (Stamberger et al., 2016; Campbell et al., 2020).

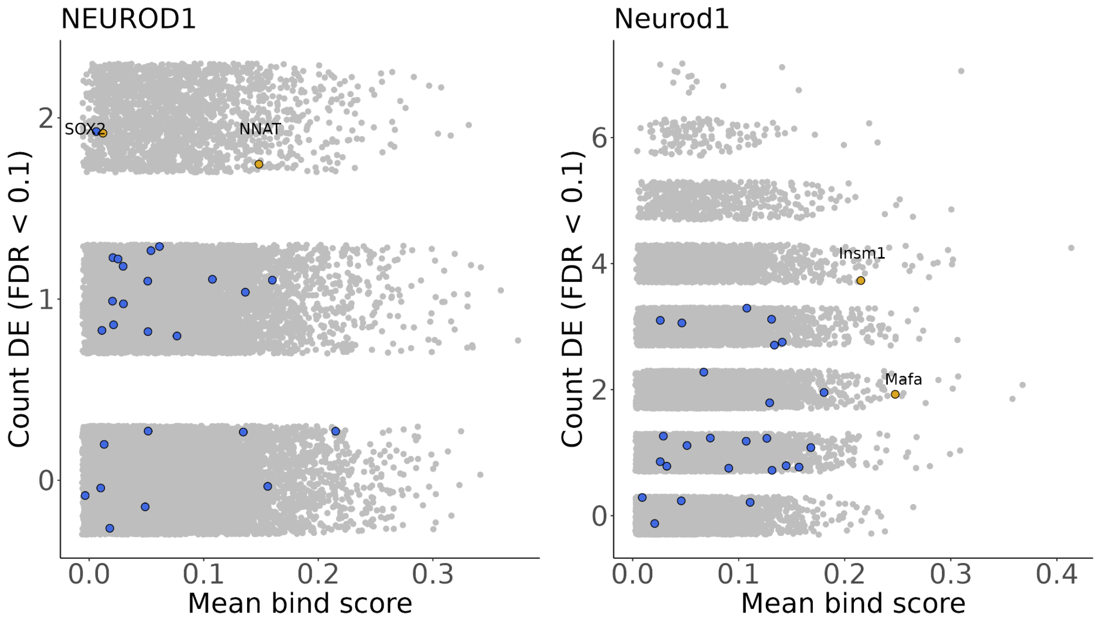

MEF2C heavily skewed towards mouse for both data types, with none of the human experiments conducted in neuronal contexts. Only a single curated target in human had a high integrated ranking, which was *MEF2C* itself (human: 3rd, mouse: 220th; Escher et al., 2011), while the top curated target in mouse was *Nr4a1* (human: 1,450th, mouse: 9th; Nagel et al., 2008). *HDAC5* (human: 158th, mouse: 96th)*, HDAC9* (human: 56th, mouse: 88th)*,* and *MEF2D* (human: 28th, mouse: 146th) were ranked highly in both species. Prior work has well established the interplay between HDACs and MEF2 TFs across multiple systems (Haberland et al., 2009), adding further support that our framework can prioritize known biology. Krupel like factors *KLF6* (human: 18th, mouse: 5th) and *KLF2* (human: 123rd, mouse: 37th) were similarly highly ranked in both species, while *KLF4* was high in mouse (human: 3,638, mouse: 65th). While absent from the curated resource, prior studies have nominated these interactions (Salma and McDermott 2012; Xu et al., 2015; Lu et al., 2021). Finally, we highlight *ARID1A* (human: 2nd, mouse: 297th) which encodes a DNA-binding SWI/SNF chromatin remodeler that was recently demonstrated to physically interact with MEF2C and regulate embryonic cardiac versus neuronal differentiation (Liu et al., 2020). Our work further suggests that MEF2C regulates expression of *ARID1A* in addition to interacting with its protein product.

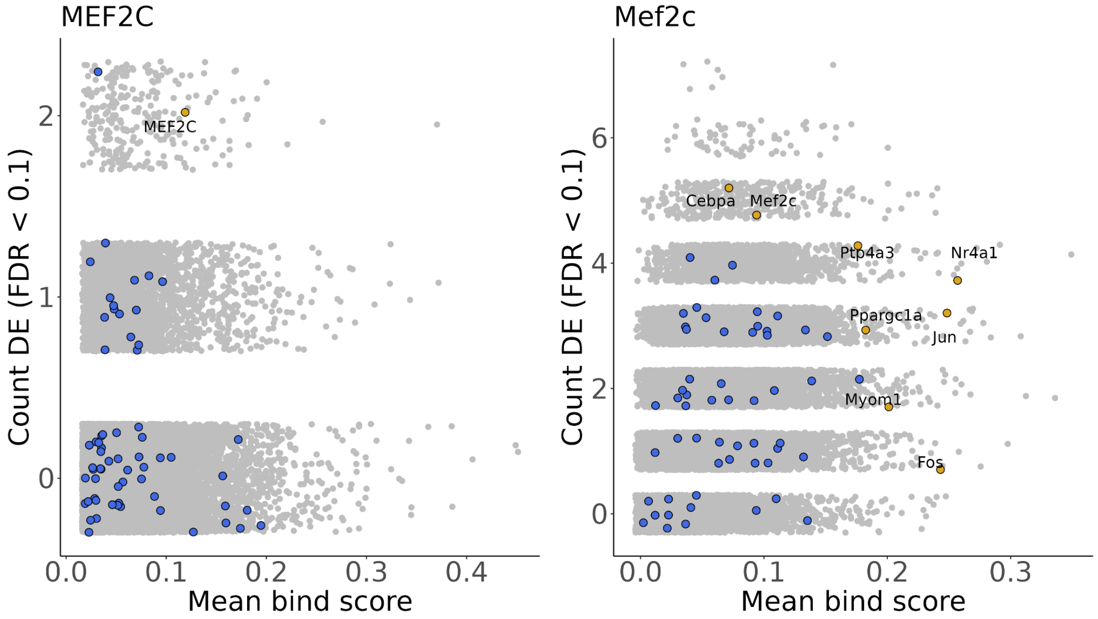

PAX6 also had more data in mouse than human, with experiments generally derived from retinal, cortical, or pancreatic tissues. After *PAX6* itself (Sun et al., 2015), high ranking curated targets included *MAB21L1* (human: 5th, mouse: 4th; Sun et al., 2015), which encodes a nucleotidyltransferase noted for its “exceptional” conservation and role in eye development (de Oliveira Mann et al., 2016). *MAB21L2* meanwhile was a curated target (Wolf et al., 2009) that ranked highly in human (3rd) but not mouse (8,825th). Retinaldehyde binding protein 1 (*RLBP1*; human: 93rd, mouse: 7th; Boppana et al., 2012) was another highly ranked and curated target with defined functionality in the eye.

Examples of top human candidates absent from the curation resource includes *EPHA3* (human: 2nd, mouse: 4,585th), which encodes an ephrin receptor kinase involved in retinotectal mapping (Lemke and Reber 2005), and *DNAJB6* (human: 1st, mouse: 2,124th), which produces a chaperone found to play a critical role in preventing neuronal protein aggregation (Thiruvalluvan et al., 2020). The lead mouse target was the biliverdin reductase-encoding *Blvra* (human: 14,434th, mouse: 2nd), with contributions from pancreatic as well as neuronal experiments. *Blvra* was also the most significant gene in the mouse binding specificity analysis (Fig. S4B). Other uncurated candidates include the neuronal marker *MAP2* (human: 117th; mouse: 66th), the poorly characterized *ZNF608/Zfp608* (human: 43rd, mouse: 219th), and *ABHD4* (human: 606th, mouse: 60th), whose lysophospholipase product was described as protective against the inappropriate delamination and migration of *Pax6*-expressing murine radial glial progenitor cells (László and Lele et al., 2020).

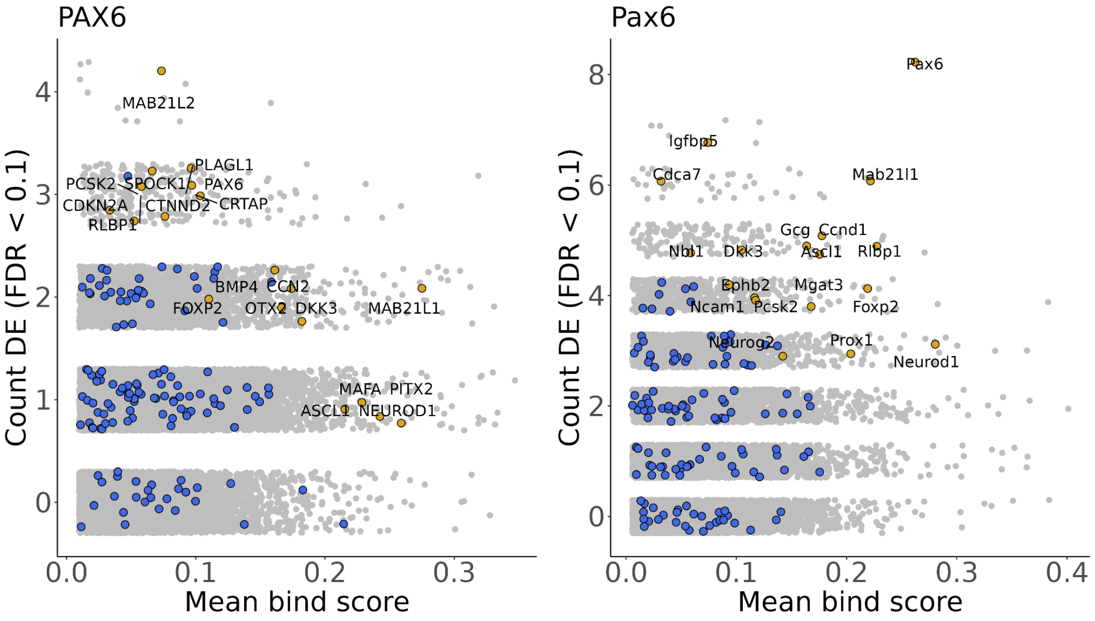

TCF4 perturbation experiments for both species covered a range of neuronal contexts, as well as experiments in blood and kidney systems. Only a single mouse Tcf4 ChIP-seq experiment was identified (a neural stem cell line; Moen et al., 2017), while the human ChIP-seq experiments tended to be from blood cancer and neuroblastoma cell lines. TCF4 was the most difficult TR to curate given the widespread prevalence of the *TCF4* symbol being used for *TCF7L2* (also known as T-Cell Factor 4), a TF commonly studied in Wnt signalling and colon cancer. A warning of this potential for confusion is also found in Papes et al., 2022. We suspected that this naming “contamination” affected other literature curated resources, and after screening all candidates only eleven *TCF4* targets were recovered.

*TCF4* (human: 7th, mouse: 1,307th) has been shown to have autoregulatory functionality in murine dendritic cell differentiation (Grajkowska et al., 2017). However, binding was low in human and not detected in the single mouse Tcf4 ChIP-seq data set, unlike for the curated target and cyclin-dependent kinase inhibitor encoding *CDKN1A* (human: 15th, mouse: 277th; Deliri et al., 2011). Lead human candidates include *DEPP1* (human: 1st, mouse; 11,563rd), an autophagy regulator gene, while the microfibril-associated *MFAP4* (human: 2nd, mouse: 13th) ranked highly in both species, as did the neurodevelopmental TF gene *ZBTB18* (human: 6th, mouse: 30th). Noting that the binding was driven by a single experiment, mouse was led by *Gpsm2* (human: 5,294th, mouse: 1st), affiliated with a human hearing phenotype (Doherty et al., 2012), and the melanocortin receptor *Mc4r* (human: 15,038th, mouse: 2nd), commonly studied in appetite control (Asai et al., 2013). Finally, *TLCD1* (human: 23rd, mouse: 3rd) was a top candidate in both species, with the sparse literature for this gene pointing to its role in membrane fluidity regulation (Ruiz et al., 2018).

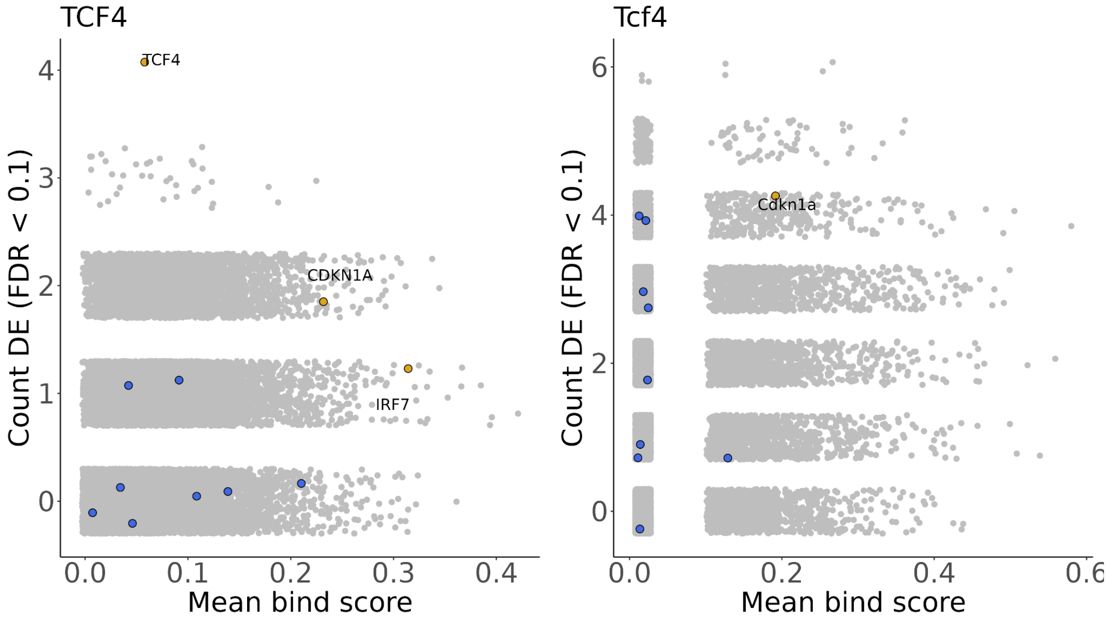

HES1 ChIP-seq data was scarce and the perturbation experiments had few DE genes (only 154 genes were DE at least once across all 9 experiments in both species), leading to less consistent aggregated evidence. Lymphotoxin beta *LTB* (human: 7th, mouse: 3,397th) was the only gene to be DE more than once in the human HES1 experiments, with binding signal in the K562 but not HepG2 or MCF7 ChIP-seq experiments. *HES1* (human: 1st, mouse: 1st) was the top curated target in both species, and has been characterized as having auto-repressive functionality (Havrda et al., 2008). *ATOH1* (human: 45th, mouse: 6,279th) was a curated target that was also the most significant gene in the HES1 specific binding analysis (Fig. 2B), but the perturbation evidence was negligible.

Of the human genes with one DE count, the mitochondrial lipid transporter-encoding *STARD7* (human: 2nd, mouse: 7,284th) had the strongest binding score, while TF gene *E2f5* (human: 1,069th, mouse: 3rd) had the strongest bind score among mouse genes with one DE count. The neurodegenerative-associated Prolifin (*Pfn1*; human: 88th, mouse: 5th; Yang et al., 2016) had the strongest binding signal in mouse — while it was not DE in the mouse studies, *PFN1* was DE once in human. Finally, *BAHCC1* (human: 29th, mouse: 292nd), which encodes a repressive histone mark reader (Fan et al., 2020), was highly bound in both species, as was the cell-cycle-associated *FBXO31* (human: 30th, mouse: 125th).

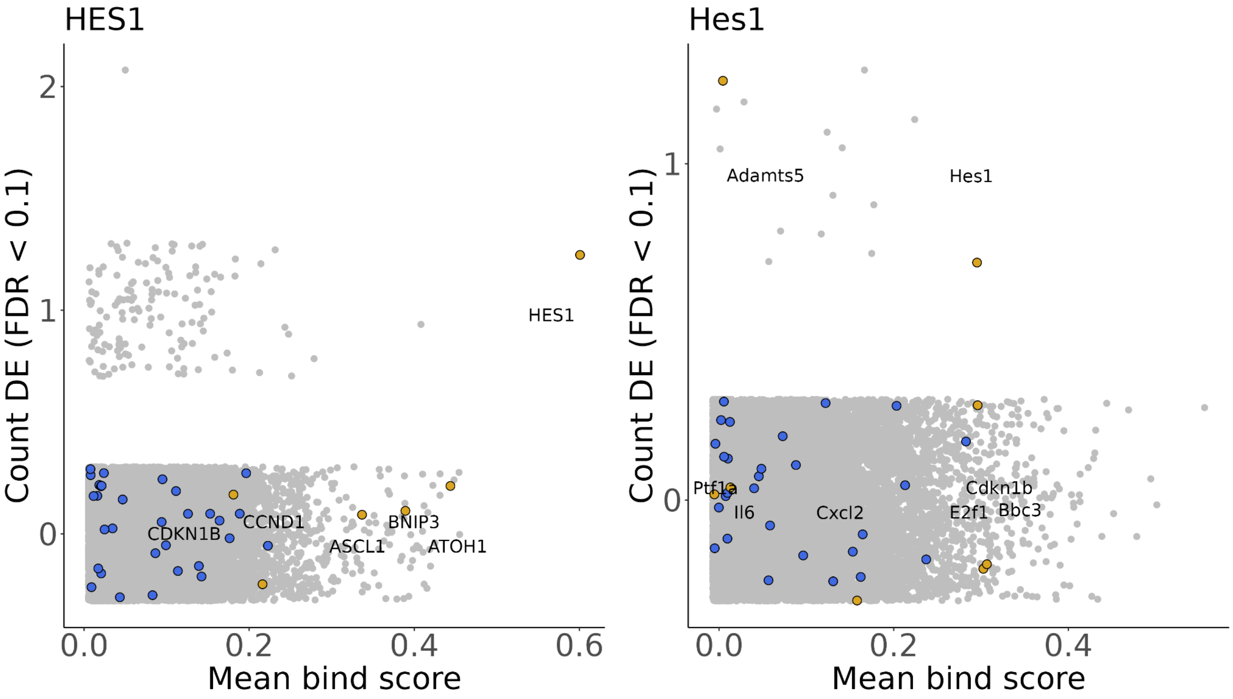
